# Earlier leaf out and loss of cold tolerance for northeastern U.S. trees in response to winter warming events and early springs

**DOI:** 10.1101/2025.05.14.653502

**Authors:** Laura Pinover, John R. Butnor, Nicholas Fisichelli, Paula Murakami, Nicole Rogers, Yong-Jiang Zhang, Jay Wason

## Abstract

- Winter warming events and early springs can advance spring phenology of trees. However, these events can also decrease cold tolerance, leaving them vulnerable to subsequent refreezing. The goal of this study was to determine how spring phenology and cold tolerance of ten tree species in the northeastern U.S. responded to warming events of different timing and duration.
- We exposed 300 tree seedlings of ten species to six different warming treatments. We assessed budburst, leaf out, vigor, and freezing damage on a weekly basis and measured the cold tolerance of species exposed to warming.
- The timing of leaf out varied among species, and generally warming treatments advanced leaf out 5 - 41 days. Leaf out was best predicted by photoperiod for deciduous trees and thermal time for evergreen trees. Cold tolerance was reduced for most species (5 - 29 °C) when exposed to only two weeks of warming, reducing their safety margin from freezing damage.
- Winter warming events and early springs will drive species-specific and foliar habit responses in phenology and cold tolerance that are likely to shape competitive forest dynamics, especially at forest ecotones.

## Introduction

Climate change is increasing the likelihood of winter warming events and early springs that impact tree phenology (Stuble et al., 2021; Zhang et al., 2022). Extreme warming events in winter can cause premature budburst of woody plants (Ladwig et al., 2019) and warming events in early spring can advance leaf out for some woody species by as much as a month (Fahey, 2016). While early budburst may increase the length of growing seasons (Morin et al., 2010), sensitive new leaves and associated decreases in cold tolerance of other tissues increases the risk of freezing damage (Vitasse et al., 2014; Baumgarten et al., 2023). However, there remains uncertainty about which tree species are most at risk and how warming events of different types, timing, and magnitude impact tree phenological responses.

Phenological responses to warming depend on the timing and severity of warming. For example, early spring warming can lead to greater advances in the timing of budburst and leaf out compared to late spring warming (Montgomery et al., 2020). Advances in the timing of budburst and leaf out can be beneficial for those plants by extending the growing season (Richardson et al., 2010). However, multiple year warming studies have shown that these advances in phenology diminish with time (Lu et al., 2025), and short events, such as 1-day experimental warming events, often have no effect on spring phenology (Marquis & Lajoie, 2023) suggesting that the duration of warming is important. Despite these patterns, many studies find different responses to warming by species or leaf functional types (Parmesan, 2007; Thackeray et al., 2016), such as angiosperms species leafing out earlier than gymnosperm species (Panchen et al., 2014). Combined, these studies suggest that there is a need for regional studies of warming effects on phenology of locally important species and leaf functional types and how they respond to warming of different durations.

Spring phenology of trees is driven by three primary cues: chilling, photoperiod, and exposure to warm temperatures (thermal time) (Flynn & Wolkovich, 2018; Satake et al., 2022). Chilling requirements, whereby exposure to sustained cold temperatures is needed before spring phenology is initiated, are common for species adapted to seasonally cold climates, assisting in protection from winter injury and are known to vary across species (Körner & Basler, 2010; Körner et al., 2016; Malyshev et al., 2023). Indeed, increased chilling can lead to earlier budburst for some temperate tree species (Nanninga et al., 2017). However, photoperiod also plays an important role in dormancy release of buds and initiating leaf out providing a seasonal cue that is independent of changing temperatures (Zohner & Renner, 2015). Finally, increased exposure to warming increases thermal time that can both advance or delay spring phenology depending on how warming interacts with species-specific chilling requirements and photoperiod (Ettinger et al., 2020; Meng et al., 2021). Thus, given the complexity of tree phenological responses to warming, and current conflicting opinions regarding the importance of these cues for predicting future forest responses to climate change (Zohner et al., 2016; Meng et al., 2021), we need to better understand the effects of chilling and photoperiod on spring phenology Trees are adapted to avoid freezing damage to tissues during winter and this degree of cold tolerance is often correlated with their distributions and likelihood of exposure to extreme temperatures (Strimbeck et al., 2007; Kreyling et al., 2015). However, warming is known to reduce cold tolerance and can leave trees susceptible to damage if cold temperatures return (Man et al., 2016). Since positive growth responses to warming are common at the cold range limit of a species (Reich et al., 2015), this can leave some species at a higher risk of cold damage than others. Cold damage can lead to restrictions in growth by reducing carbon stores and reducing leaf area for photosynthesis (Schaberg et al., 2011; Bhattacharya, 2022). Although there are examples of freezing damage following early warming events (Frelich et al., 2021), after exposure to warming some tree species can regain their cold tolerance (Man et al., 2014; Chang et al., 2021). Furthermore, compared to evergreen trees, deciduous trees completely rely on new leaves each season. Therefore deciduous species may be adapted to generally leaf out earlier, as long as reliable cues that winter is over have been received (adequate photoperiod or chilling).

As winter warming events and early springs become more common with climate change, the likelihood of trees being exposed to refreezing events after warm periods is increasing. However, we know very little about how these warming events impact cold tolerance for many species and the degree to which it leaves them vulnerable to subsequent refreezing.

In this study, we used experimental warming treatments to determine how winter warming events and early springs impact the phenology and cold tolerance of containerized seedlings of 10 tree species native to the northeastern USA. We focus on seedlings because they can be the most vulnerable growth stage and their growth and survival determines future forest dynamics (Fenner, 1987; Winkler et al., 2024). We tested the extent to which the timing of leaf out across species’ leaf functional types (deciduous vs. evergreen) responded to warming events of different lengths and at different times of the year. We also tested how warming events at different times of year impacted cold tolerance and subsequent damage from exposure to refreezing. We hypothesized that: i) in response to warming events, deciduous species would advance leaf out earlier than evergreen species; ii) the effects of thermal time on leaf out will depend on time of year and species-level chilling and photoperiod requirements iii) the species that exhibited the strongest leaf out responses to warming and had the most southern ranges would also experience the largest loss of cold tolerance.

## Materials and Methods

### Study species and site

We conducted this study at the University of Maine, Roger Clapp Greenhouse Complex in Orono, ME (44.8834° N, −68.6708° W; 36-m above sea level). We selected ten tree species common to the Northeastern United States (five deciduous: *Acer rubrum* L.*, Acer saccharum* Marsh.*, Betula papyrifera* Marsh.*, Castanea dentata* (Marshall) Borkh.*, Quercus rubra* L. and five evergreen*: Abies balsamea* (L.) Mill.*, Pinus resinosa* Ait.*, Pinus strobus* L.*, Picea rubens* Sarg., and *Juniperus virginiana* L. (Table 1). These species represent a range of distributions and functional types that may relate to differences in phenology and cold tolerance. We sourced seedlings of one to three years of age for each species as bare-root stock from two nurseries similar in climate and latitude to Maine (Table 1) besides *Castanea dentata* (Marshall) Borkh. which were pure *Castanea dentata* (Marshall) Borkh. and grown from seed sourced from two orchards in central Maine. In early May 2023, we planted 21 individuals per species each into 7.5-L containers filled with a soilless potting mix (Jolly Gardener CF) amended with slow-release fertilizer (Osmocote Plus 15-9-12 N-P-K; 3.25 g L^-1^). Each seedling was assigned to one of three experimental blocks based on height and randomly assigned to one of seven treatment groups (including 2 control groups) within that block yielding one species per treatment group per block. The seedlings were grown outdoors in mostly full-sun conditions and we used an overhead irrigation system to water seedlings to saturation twice a week until the end of September 2023. For overwintering, straw was packed between containers to insulate roots. We measured height, diameter, and vigor immediately after seedlings were planted, at the end of the growing season in 2023 and 2024, and at the end of the warming treatments. We determined height by measuring the length to the longest shoot and diameter as the average of two perpendicular measurements 2-cm above the media surface. We also classified seedlings into one of three vigor categories from healthy to dead.

**Table 1.** Seedling species, source, range in relation to Maine, age, height and diameter for the spring 2023 planting. Species are split up by deciduous and evergreen species.

| Species | Source | *Range | Age | Height (cm) | Diam (mm) |
| --- | --- | --- | --- | --- | --- |
| *Acer rubrum* | Michigan | Northern/Central | 1 | 33.4 ± 15.7 | 7.3 ± 2.9 |
| *Acer saccharum* | Michigan | Northern | 1 | 33 ± 7.5 | 4.4 ± 1.5 |
| *Betula papyrifera* | Michigan | Southern | 1 | 45.6 ± 19 | 4 ± 1.9 |
| *Castanea dentata* | Maine | Northern | 1 | 16.4 ± 13.5 | 2.5 ± 1.3 |
| *Quercus rubra* | New Hampshire | Northern | 2 | 16.5 ± 11 | 3.7 ± 2.2 |
| *Abies balsamea* | Michigan | Southern | 3 | 25.7 ± 8.9 | 5.9 ± 2.8 |
| *Juniperus virginiana* | New Hampshire | Northern | 2 | 17.7 ± 9.5 | 2.9 ± 1.5 |
| *Picea rubens* | New Hampshire | Southern | 3 | 24.4 ± 11 | 4.1 ± 3.5 |
| *Pinus resinosa* | New Hampshire | Southern | 3 | 30.3 ± 10.7 | 6.2 ± 2.9 |
| *Pinus strobus* | Michigan | Northern/Central | 3 | 30 ± 16.9 | 4.5 ± 1.8 |

\*Maine, U.S. in location to native range

We monitored hourly temperature and humidity outside and inside the greenhouse using Onset HOBO External Temperature/RH Sensor Data Loggers (two outside and two in the greenhouse where we applied warming treatments, see next paragraph). We also monitored hourly soil temperature using 14 iButton data loggers (Thermochron model DS1921G, resolution 0.5°C; Maxim Integrated Products, Inc., Sunnyvale, California) sealed in latex balloons and each buried 10-cm below the media surface in a separate 7.5-L containers without seedlings. To track how soil temperature responded to treatments, each container with an iButton was then randomly assigned to one of each of our 6 treatments yielding two iButtons per warming treatment and four outdoor controls. Overall, we found there was approximately a two-day delay in soil temperature in order to reach similar air temperatures when seedlings were moved into the greenhouse (data not shown) and we therefore focused our analysis on air temperatures.

### Warming treatments

In order to determine how spring phenology responded to warming, we simulated warming events by moving seedlings inside a greenhouse at different times of year and for different durations (Table 2). The greenhouse experienced natural light. Although our control of temperature was limited, the average daily temperature in the greenhouse during our study was 16.3°C ± 2.3°C (Fig. S1). We tested five different warming treatments (Table 2): 14-day warming events in late February (F14) and late March (M14), a repeated 7-day warming event in late March and mid-April separated by one week outdoors (M7R), an extended early spring event in early April (ES), and an event that occurred at the beginning of March where each seedling was inside the greenhouse until it reached leaf out (M2LO).

**Table 2.**
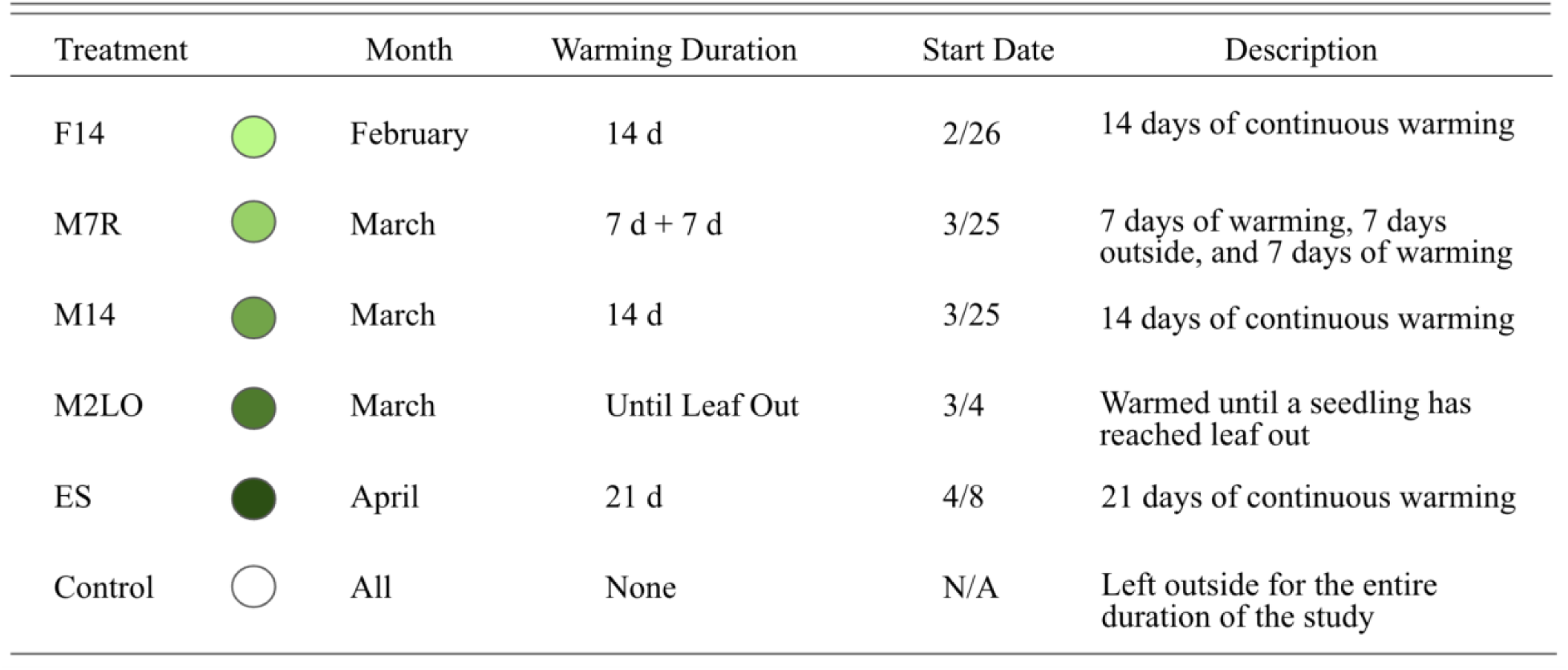
Timeline of when seedlings experienced warming treatments. Each treatment includes the month the warming took place, duration, start date, and further description of the warming treatment. Colors represented for each treatment are consistent throughout all figures.

| Treatment |  | Month | Warming Duration | Start Date | Description |
| --- | --- | --- | --- | --- | --- |
| F14 |  | February | 14 d | 2/26 | 14 days of continuous warming |
| M7R |  | March | 7 d + 7 d | 3/25 | 7 days of warming, 7 days outside, and 7 days of warming |
| M14 |  | March | 14 d | 3/25 | 14 days of continuous warming |
| M2LO |  | March | Until Leaf Out | 3/4 | Warmed until a seedling has reached leaf out |
| ES |  | April | 21 d | 4/8 | 21 days of continuous warming |
| Control |  | All | None | N/A | Left outside for the entire duration of the study |

### Monitoring Phenology

We quantified the phenophases for each seedling weekly from February 14, 2024 through June 16, 2024. We built custom species level phenology guides based on those available for the same or related species from the National Phenology Network and The University of Vermont and using our own photos from this study (Rosemartin et al., 2018; Skinner & Parker, 1994). Our phenology guides included up to eight phases for each species from bud dormancy, through leaf out (examples in Fig. S2), to fully unfolded leaves. When scoring phenophases, each tree was scored a majority phenophase (i.e. the majority phase of all buds) and a maximum phenophase (i.e., the maximum phase of any bud). We analyzed our results both ways and found no substantial differences in results or overall conclusions. Therefore, and based on previous studies, we decided to focus our results on the majority phase (Miller-Rushing et al., 2008).

Additionally, we focused on the leaf out phenophase because it is the most sensitive phenological stage (Vitasse et al., 2014). We assessed tissue damage as seedlings were brought back outside after warming by visually estimating the percentage of the new growth with damage.

### Cold Tolerance Sampling

To determine how warming impacted the cold tolerance of each species, we measured the cold tolerance of each species on March 8, April 11, and April 22, 2024. During each sampling event, we collected tissue from two seedlings per species (to limit the amount of sampling damage to seedlings in our study): one control seedling (no exposure to warming) and one warmed seedling that had been exposed to two weeks of warming in the greenhouse from one of our treatments.

For each seedling, we removed a 6-8 cm section of stem tissue (deciduous species) or approximately 5-20 needles (evergreen species). Stem tissue was not used for evergreen species due to the limited amount of tissue available. Unfortunately, we could not sample from *Castanea dentata* (Marshall) Borkh. and *Quercus rubra* L. because of their small size and limited available tissue. After the samples were collected, we cut the stem tissue (excluding buds) or needles into 3-mm pieces and placed five pieces originating from the same species and treatment (control vs. warmed) into three wells within well plates. Therefore, for each sampling round, we had three to five measurement replicates per species per treatment per round. Immediately after the samples were put into the wells, the plates were wrapped with parafilm and put into coolers with ice, which were then shipped overnight to USDA Forest Service, NRS Laboratory in South Burlington, VT and refrigerated until processing within 24 hours of sample collection. Samples were randomly assigned to be exposed to a stepwise freezing processing involving +4°C, −10°C, - 20°C, −40°C, or −80°C (with additional temperatures at −5°C, −15°C, and −25°C during the third sampling round). In order to measure cold tolerance, we followed (Butnor et al., 2024). The samples were put into solution and we measured initial conductivity post thawing and final conductivity after heat exposure, which induced electrolyte leakage.

Historical daily minimum temperatures were calculated using data from the NOAA’s National Centers for Environmental Information climate data records.

### Statistical Analysis

Leaf out is a critical phenophase of spring phenology and is readily identifiable (Fig. S2). Therefore, we summarized our phenological data to a binary variable indicating if the seedling had the majority of its buds reach the phenophase leaf out or not. While our focus for our study focuses on the leaf out phenophase, we also repeated our analysis for majority budburst and found that results were very similar (see Supplemental Figures).

To test the estimated date of leaf out for each species and treatment, we fit separate generalized linear models for each species and treatment combination with leaf out as a binomial response variable and DOY as the predictor. We then used those models to predict the DOY when 50% majority leaf out occurred and calculated 95% confidence intervals for those estimates. In order to determine if there were consistent effects of warming treatment on leaf out date across species, for each species we calculated how the DOY for majority leaf out in each treatment differed from controls for that species. We also used those values to calculate an average difference from controls across species. In both the species-level and across species summaries for the change in leaf out relative to controls, if 95% confidence intervals did not overlap zero, we took those to be significantly different from controls. We also used ANOVA to test if effects were significantly different among warming treatments.

Next, we modeled how thermal time, chilling, and photoperiod could predict the timing of leaf out among all treatments for each species. To quantify thermal time and chilling units, we calculated accumulated growing degree days (GDD) and accumulated chilling days for each treatment using a combination of the outdoor or greenhouse data logger data depending on when that treatment was exposed to each condition. GDD for a given day was calculated as:
(T_max_ + T_min_)/2-T_base_
where T_max_ is the maximum daily temperature and T_min_ is the minimum daily temperature. The base temperature (T_base_) was set to +5°C. Chilling days were calculated as the number of days ≤ T_base_ since November 1, 2023. T_base_ was set to +5°C (Cannell & Smith, 1983). The suncalc package was used to estimate photoperiod for each day (Thieurmel & Elmarhraoui, 2022).

Unfortunately, collinearity among GDD, photoperiod, and chilling days (Fig. S3) resulted in high variance inflation factors for our models. Therefore, we ran two separate modeling approaches to examine how GDD, which is known to impact spring phenology, interacted with either photoperiod or chilling days separately. First, to test how leaf out responded to GDD and photoperiod, we developed species-level generalized linear models predicting leaf out as a function of GDD, photoperiod, and their interaction. The interaction was removed if it was not significant. GDD and photoperiod were standardized prior to model fitting (mean = 0, SD = 1) and model coefficients are reported as effect sizes so they are comparable units of standard deviations. Next repeated this process replacing photoperiod with chilling days.

To examine how cold tolerance responded to warming at different times of year, we modeled relative electrolyte leakage for each species, treatment, and round combination as a non-linear dose-response function with temperature as the predictor using the drm function in the drc package in R (Butnor et al., 2024; Ritz et al., 2015). From that model, we extracted the lethal temperature reaching 50% of maximum electrolyte leakage (LT_50_). Differences between LT_50_ of control and warmed seedlings were considered significant if the 95% confidence intervals did not overlap.

Finally, to determine if there were significant differences in height growth among warming treatment or species, we calculated height growth at the end of the growing season following our warming treatments as a proportion of starting height that spring. We used multiple linear regression models predicting relative height growth as a function of species and treatment and their interaction (interaction removed if not significant). Significant differences among predictor groups were determined using Tukey’s Honest Significant Difference (Tukey, 1953). All statistical analysis and graphing was performed in R (ver. 4.4.2, R Core Team 2024) using RStudio (Posit Team, 2024).

## Results

### Leaf out phenology

For control seedlings, we found a 43 d difference between the earliest species to leaf out (*Betula papyrifera*: DOY 123) and the latest (*Pinus resinosa*: DOY 166; Fig. 1). With the exception of *Quercus rubra* (147 d), deciduous species leafed out earlier than evergreen species.

**Fig. 1.**
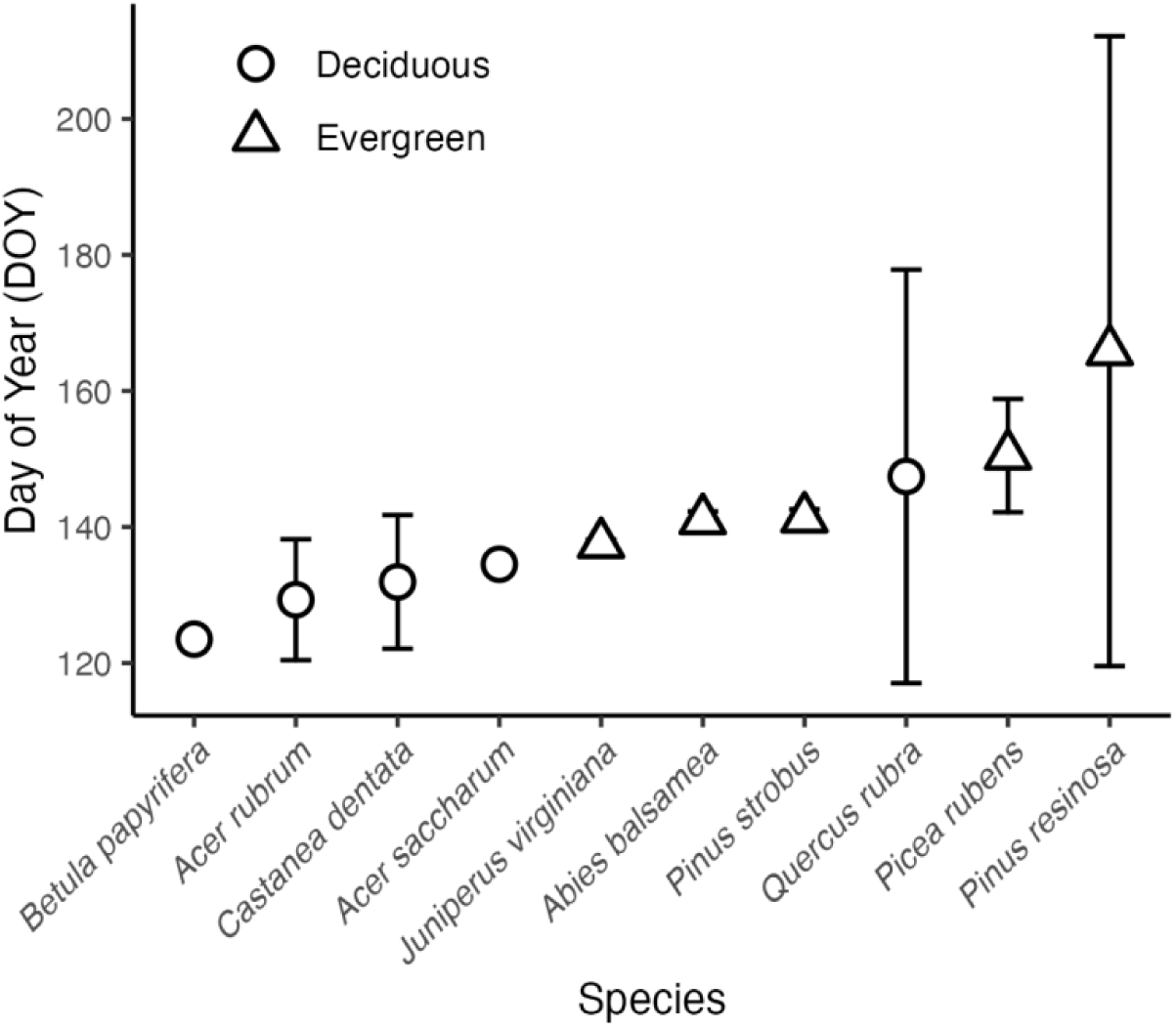
Day of year when majority leaf out appeared for our control study species represented by 95% confidence intervals. Circles indicate deciduous species and triangles indicate evergreen species. Species with no visible error bars had little or no variation in leaf out date that was detectable by our weekly resolution.

When we averaged across species, we found that all but the M7R treatment significantly advanced leaf out compared to controls (Fig. 2). On average, leaf out was advanced the most in M2LO (−20.7 d) and ES (−16.2 d) treatments. Although there was substantial variability, fourteen days of warming in March (M14) advanced leaf out −9.1 d compared to only −5.5 d for the same length of warming in February. Interestingly, the repeated warming treatments separated by one week of control temperatures (M7R) did not significantly advance leaf out whereas two-weeks of continuous warming without the break did advance leaf out by an average of −9.1 d (M14).

**Fig. 2.**
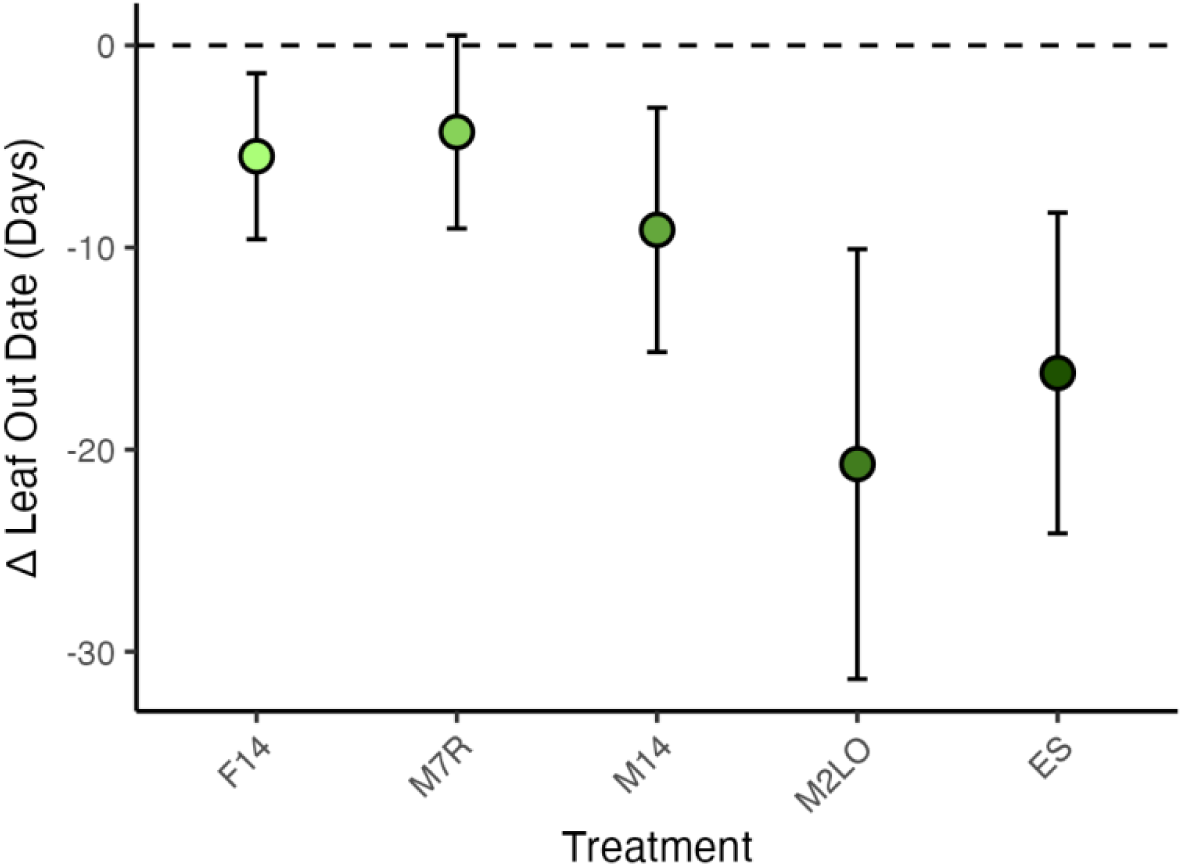
The mean difference between treatment leaf out date from control leaf out date represented by 95% confidence intervals. Treatment is significant if the confidence intervals fall below 0.

Next we examined species level differences in leaf out date by treatment and we found earlier leaf out among most species in the ES treatment for *Betula papyrifera* (−17.3 d; Fig. **3A**), *Acer saccharum* (−18 d; Fig. **3G**), *Juniperus virginiana* (−16.3 d; Fig. **3B**), *Pinus strobus* (−28 d; Fig. **3F**), and *Quercus rubra* (−37.9 d; Fig. **3I**). The M2LO treatment also yielded significant advances in leaf out for four of the five evergreen species: *Juniperus virginiana* (−39.7 d; Fig. **3B**), *Abies balsamea* (−41.7 d; Fig. **3D**), *Pinus strobus* (−38.8 d; Fig. **3F**), and *Picea rubens* (−34 d; Fig. **3H**). However, the other three treatments, F14, M14, and M7R, showed minor shifts compared to controls when analyzed at the species level. *Pinus strobus* (Fig. **3F**) significantly advanced leaf out under all warming treatments compared to controls. In contrast, *Pinus resinosa* (Fig. **3J**) was the only species that had low vigor across all treatments (including controls) and showed no significant change in leaf out.

**Fig. 3.**
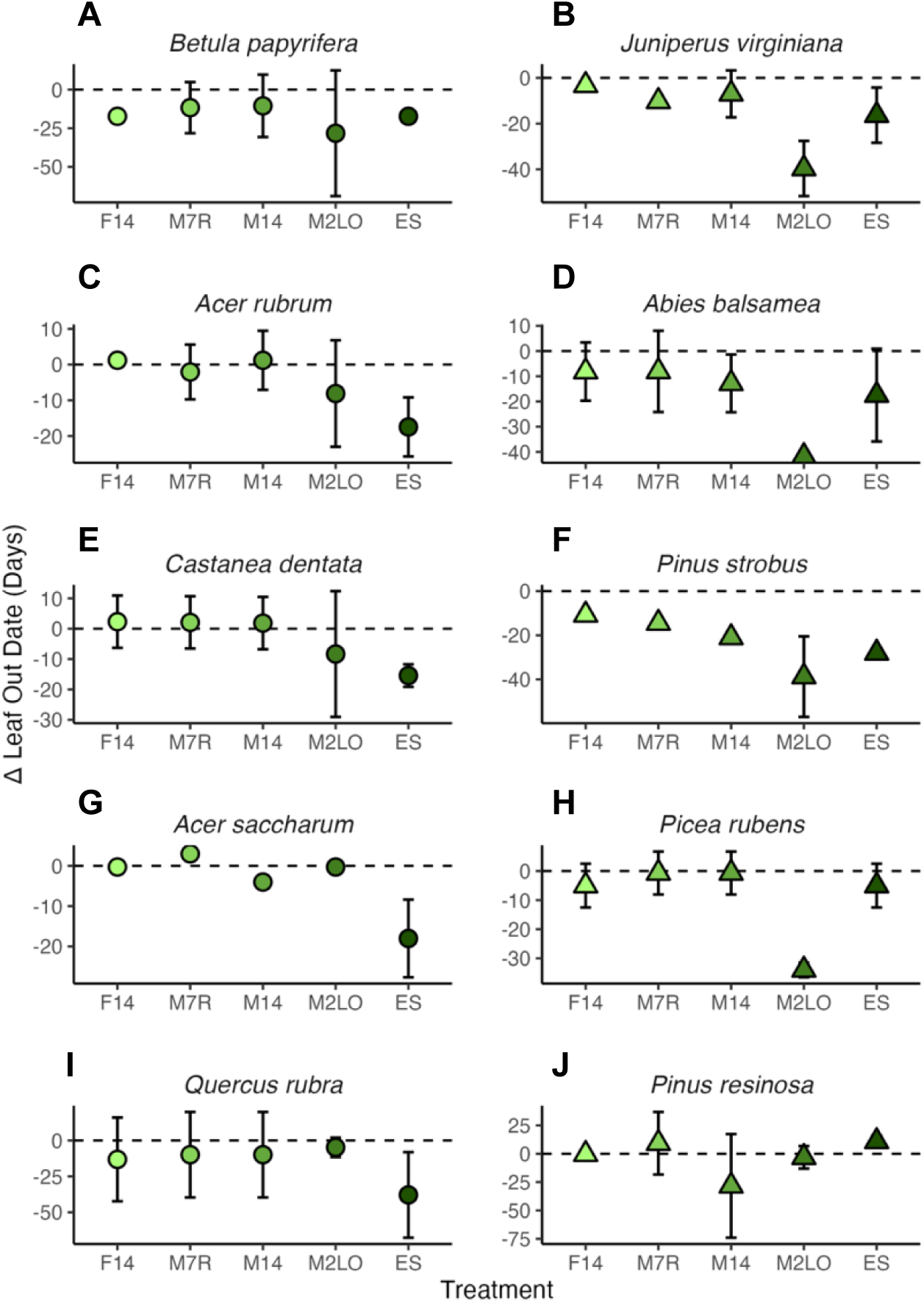
95% confidence intervals for the difference in leaf out day of year (DOY) among each species and each warming treatment and control. Each panel represents a species. Warming treatments are deemed significant if the 95% confidence intervals fall below 0. Species with each column are ordered by the leaf out date for our controls, starting with the species to leaf out the earliest to the species to leaf out the latest. Shape means with no visible error bars, have either error bars behind the shape or no variation in the mean.

### Predicting leaf out with growing degree days, photoperiod, and chilling days

We next tested how GDD and photoperiod as well as GDD and chilling days predicted leaf out. Both models explained a similar amount of variance (mean R^2^ = 0.77) and we found overall similar effect sizes in our GDD and photoperiod models (Fig. 4) and in our GDD and chilling models (Fig. 5). We found that leaf out of deciduous species tended to respond more strongly to photoperiod than GDD with longer photoperiod generally increasing the probability of leaf out (Fig. **4A**). In contrast, leaf out of evergreen species responded more strongly to GDD than photoperiod (Fig. **4B**). For example, for *Juniperus virginiana*, *Abies balsamea*, *Picea rubens,* and *Pinus strobus* the effect size for GDD was more than 2.5 times greater than for photoperiod. Leaf out of the other deciduous species, *Pinus resinosa*, also responded strongly to GDD, but there was a significant negative interaction with photoperiod suggesting that the effect of GDD decreased at longer photoperiods. In contrast, leaf out of the deciduous species *Acer rubrum*, *Castanea dentata*, *Acer saccharum*, and *Quercus rubra* was more strongly predicted by photoperiod than GDD and for *Acer saccharum,* this effect was stronger at longer photoperiods. However, for *Betula papyrifera*, the earliest species to leaf out in our study, leaf out was best predicted by GDD with a negative interaction with photoperiod suggesting that the effect of GDD on leaf out became weaker at longer photoperiods. Results from our models with GDD and chilling days were slightly different with GDD being the dominant positive predictor for all species (Fig. 5). Chilling days had a small positive or no effect on leaf out and there were only minor positive (*Pinus strobus*) or negative (*Picea rubens*, *Pinus resinosa*, and *Castanea dentata*) interactions between GDD and chilling days.

**Fig. 4.**
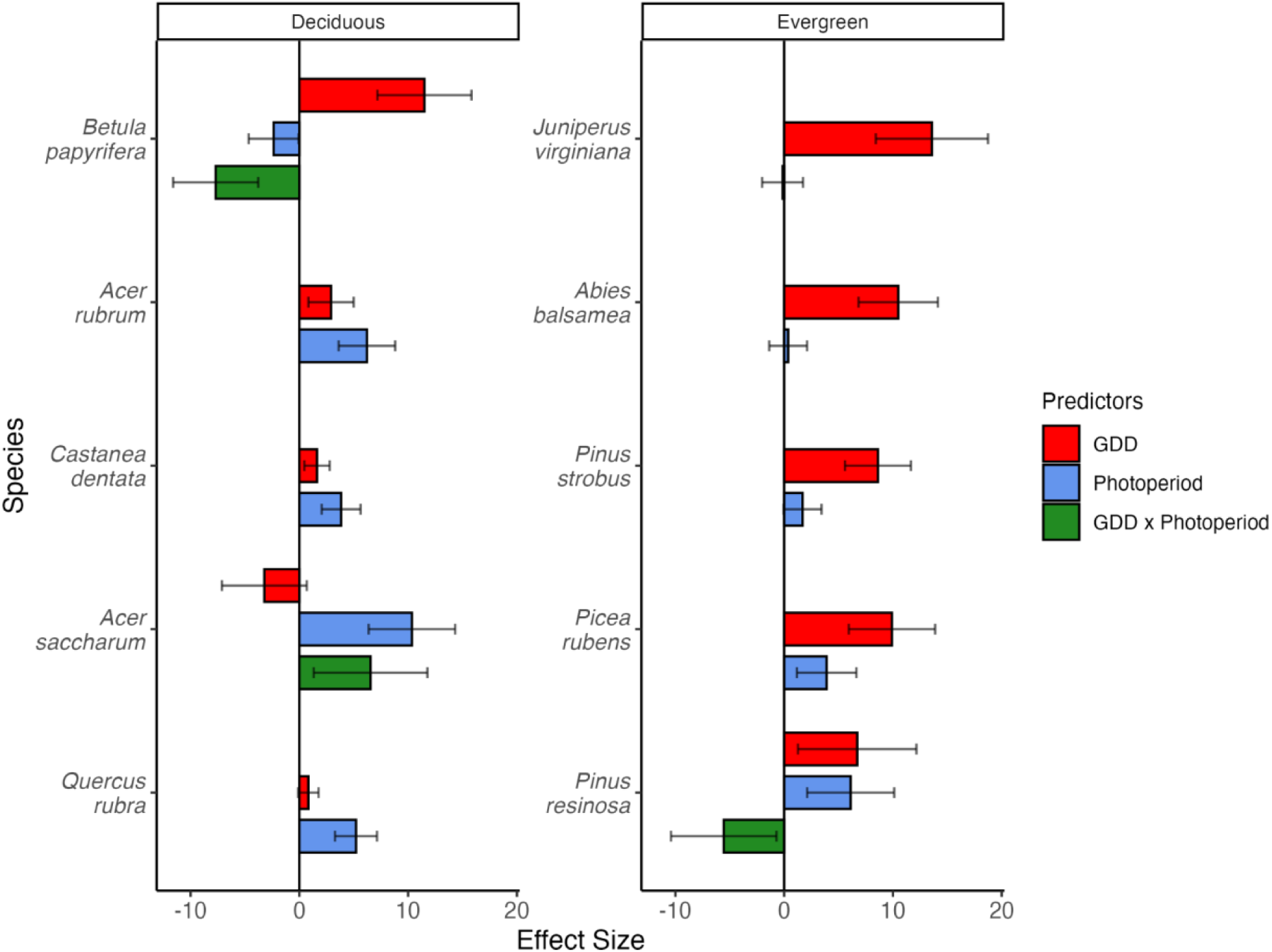
Effect sizes for leaf out predictors Growing Degree Days (GDD), photoperiod, and the interaction between GDD and photoperiod for all study species from generalized linear mixed effects models. Individual models were created for each species. Columns are split by deciduous and evergreen species. Species with all three predictors present included the interaction between GDD and photoperiod in the model, whereas species with two predictors had the interaction removed from their models.

**Fig. 5.**
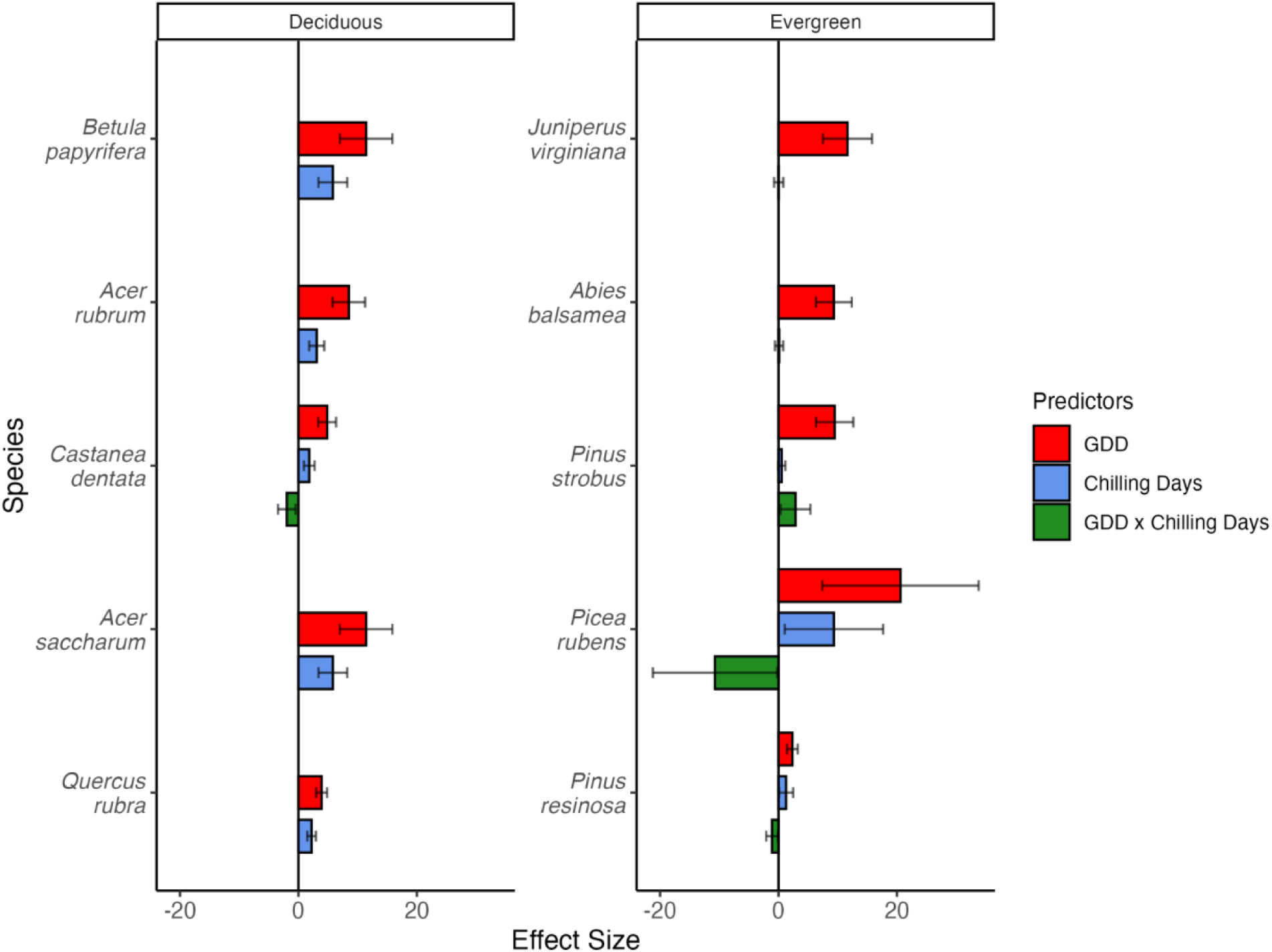
Effect sizes for leaf out predictors Growing Degree Days (GDD), Chilling Days, and the interaction between GDD and Chilling Days for all study species from generalized linear mixed effects models. Individual models were created for each species. Columns are split by deciduous and evergreen species. Species with all three predictors present included the interaction between GDD and Chilling Days in the model, whereas species with two predictors had the interaction removed from their models.

### Cold tolerance

Next, we assessed how two weeks of warming at three different times throughout the study impacted cold tolerance for eight of our ten species. Overall, for control seedlings we found that cold tolerance was lowest in the first or first two sampling events and, except for *Pinus resinosa*, all species began to lose cold tolerance by the third event (Fig. 6). In the first sampling event, we found significant reductions in cold tolerance for warmed evergreen species relative to controls of *Juniperus virginiana* (−21.9°C; Fig. **6D**), *Pinus strobus* (−14.4°C; Fig. **6F**), *Picea rubens* (- 10.4°C; Fig. **6G**), and *Pinus resinosa* (−13.6 °C; Fig. **6H**) which brought them closer to historical daily minimum temperatures. In contrast, cold tolerance of *Abies balsamea* (Fig. **6E**), *Betula papyrifera* (Fig. **6A**), *Acer saccharum* (Fig. **6C**), and *Acer rubrum* (Fig. **6B**) did not change after exposure to the first warming treatment. However, by the second sampling event, *Betula papyrifera* (−29.8°C; Fig. **6A**), *Pinus resinosa* (−12.8°C; Fig. **6H**), and *Pinus strobus* (−22°C; Fig. **6F**) showed significant reductions in cold tolerance in response to warming, whereas the other species exhibited no change besides slight reductions in *Abies balsamea* (−5.6°C; Fig. **6E**) and *Juniperus virginiana* (−8.3°C; Fig. **6D**). By the third sampling event, as most controls had also lost cold tolerance, only *Betula papyrifera* (−12.9°C; Fig. **6A**)*, Pinus resinosa* (−17.5°C; Fig. **6H**), *Pinus strobus* (−8.2°C; Fig. **6F**), and *Abies balsamea* (−5.1°C; Fig. **6E**) still responded to the warming treatment by losing cold tolerance.

**Fig. 6.**
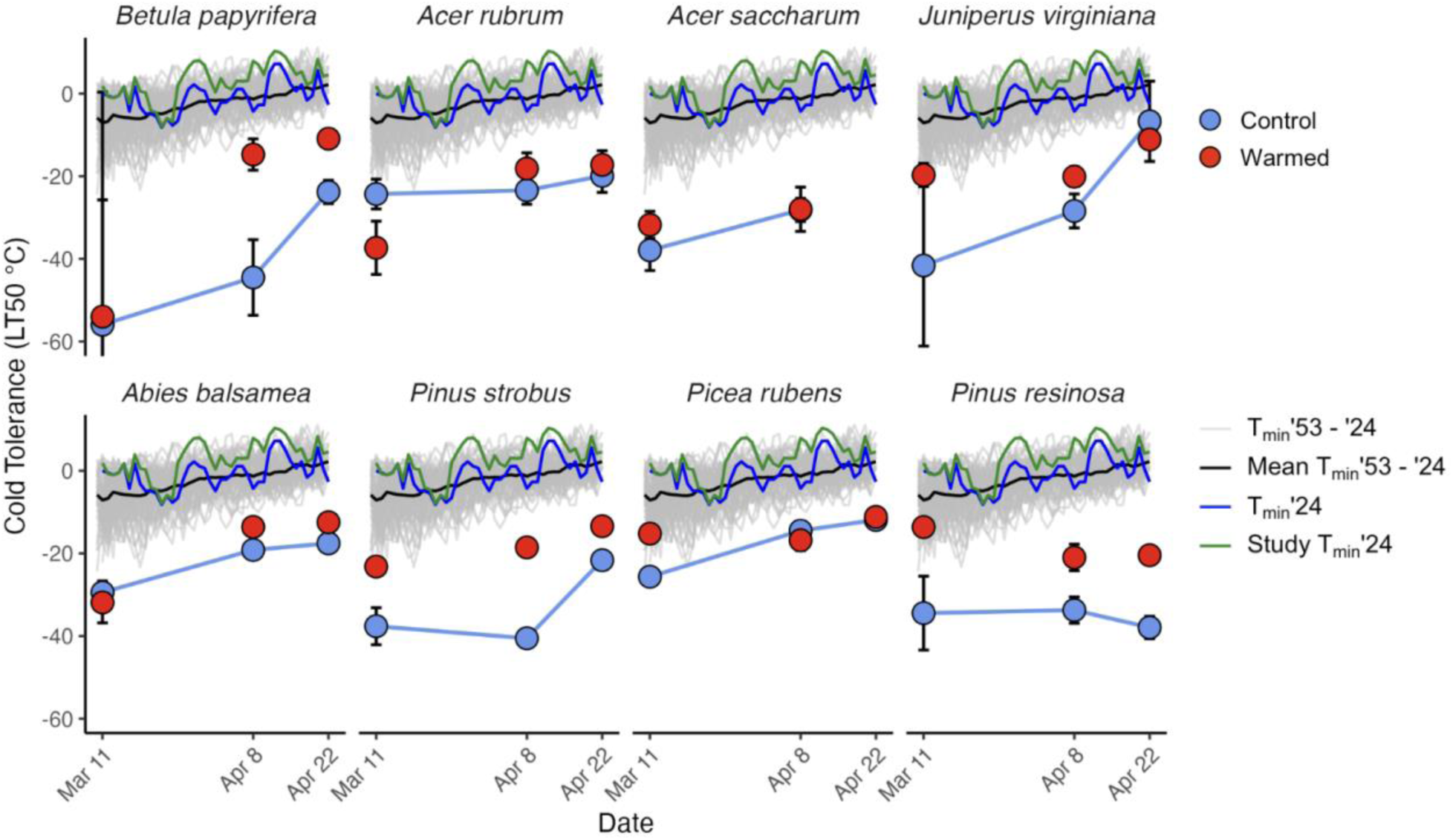
Mean lethal temperature values where 50% of cells experienced leakage for control and warmed treatments. Error bars indicate 95% confidence intervals. Date listed was the sampling date after two weeks of warming. Species ordered by average control leaf out date with the first species, *Betula papyrifera* (**A**) leafing out the earliest and *Pinus resinosa* **(H)** leafing out the latest. The black line represents the overall mean daily minimum temperature (1953 to 2025) from Bangor, Maine, while the grey lines represent daily mean minimum temperatures for individual years. The green line represents the mean daily minimum temperature from 2024 based on the Bangor Maine climate data and the blue line represents the mean daily minimum temperature from 2024 based on our study climate sensors.

### Damage from refreezing and impacts on growth

Spring 2024 was the 15^th^ warmest spring on record from Bangor, Maine climate data and we found that temperatures outside the greenhouse did not go below the cold tolerance of even warmed seedlings. Therefore, with these warm temperatures, the opportunity for re-freezing and tissue damage to occur among our study seedlings was minimal. However, we still observed damage on 28 individual seedlings, with damage primarily on *Betula papyrifera* and *Juniperus virginiana.* The majority of damage across all species was less than 10% of new growth and we found that there was no effect of this damage on growth or survival. Warming treatments did not significantly affect relative height (Fig. S4) or relative diameter growth (Fig. S6). However, there were significant differences among species for both relative height (species main effect p-value < 0.001; Fig. **5**) and relative diameter growth (Fig. S7).

## Discussion

Our results provide new information around complex spring phenology and cold tolerance patterns of NE tree species in response to warming. We revealed high variation among NE tree species in their phenological sensitivity to warming; spring phenology of leaf out for many tree species advanced 4.3 - 20.7 days in response to winter warming events and early springs.

However, the degree of phenological sensitivity to warming at different lengths at different times of the year varied by species in ways that we find may also be attributed to different dominance among environmental cues (Körner & Basler, 2010). Regardless of the drivers of spring phenology, we also found some evidence to suggest that spring phenology and cold tolerance responses to warming depended on functional type (deciduous angiosperm vs. evergreen gymnosperm) and location within a species range (Richardson et al., 2013; Zettlemoyer et al., 2019).

### Warming impacts on timing of leaf out

Overall advances in spring phenology were stronger for seedlings that experienced longer warming treatments than shorter warming treatments (Fig. 2 and Fig. S8). Similarly, in other experimental studies, longer warming contributed to the accumulation of growing degrees and led to larger advancements in spring phenology (Marquis & Lajoie, 2024). The average daily temperature in the greenhouse during our study was 16.3°C ± 2.3°C. Since the accumulation of GDD can happen rapidly with intense warming or slowly with moderate warming, future research could test the extent to which our patterns would persist in more moderate warming treatments of longer duration. It is worth noting that there may be differences in belowground phenology among seedlings planted in natural settings than compared to our study. Our data suggest that root systems reached similarly warm temperatures as the air just two days after being brought into the greenhouse. Thus, our study included belowground warming, and research has shown that phenological mismatches can occur between aboveground and belowground phenology (Liu et al., 2022). However, temperature may not be as important as other factors for belowground phenology (Abramoff & Finzi, 2015) suggesting that although responses may be slower if soils were not warmed, we may still expect the overall pattern to be similar. Regardless, future work could test how differential warming of air and soil drives spring phenology.

Generally, deciduous angiosperms were the most sensitive to the ES warming treatment. This finding could be linked to deciduous angiosperms leafing out before evergreen gymnosperms in our study (Fig. 2) and others (Panchen et al., 2014). The need for deciduous species to produce new leaves early to maximize carbon gain that season, is a fundamental tradeoff that may be less important for evergreen species (Michelot et al., 2012). Additionally, other studies have found that species with diffuse porous wood anatomy generally leaf out earlier than ring porous species (Savage et al., 2022), which may explain why in our study the last deciduous species to leaf out was ring porous (*Quercus rubra*).

### Environmental drivers of leaf out timing

We also found that, in addition to the effects of warming, photoperiod and chilling had larger effect sizes on deciduous species than evergreen species. For the deciduous species in this study, the limited phenological response to warming in winter and the need for photoperiod or chilling requirements to be met before responding to warming may be an adaptation to time leaf out correctly (Meng et al., 2021). For deciduous species producing a new full set of leaves each year, this adaptation can help protect newly-developed leaves from damage (Körner & Basler, 2010), particularly since they tend to leaf out earlier than evergreen species already. *Betula papyrifera* was the exception to this trend and leafed out early in response to some warming treatments that other deciduous species did not respond to. *Betula papyrifera* did also experience instances of damage to some leaves (discussed more later) but, this fast growing pioneer species was able to resprout effectively suggesting that its life history strategy may be riskier and allow it to take advantage of early springs (Asse et al., 2018). Overall, these results support other work that highlights the importance of GDD for leaf out but they point to an important need for future studies to separate the effects of chilling and photoperiod as limiting factors for spring phenology.

In contrast, evergreen species responded the most to the M2LO warming treatment, and our models suggest that they mostly do not have photoperiod or chilling limitations and that leaf out is mainly driven by GDD. The geographic range of evergreen species in this study may at least partly explain this difference from the deciduous species. Among our northern range evergreen species (*Abies balsamea*, *Picea rubens*, and *Pinus resinosa*), most had large effect sizes for GDD. These results match recent studies suggesting that, particularly more boreal type and northerly distributed species may advance spring phenology in response to warming (Montgomery et al., 2020; Frelich et al., 2021). Boreal species tend to have lower chilling and photoperiod requirements, if any at all, making them more susceptible to warming (Zohner et al., 2016). Combined, these results suggest that chilling and photoperiod requirements had already been met, the warming treatments were strong enough to overcome chilling and photoperiod requirements, or chilling and photoperiod requirements may be weak or absent (Montgomery et al., 2020; Pan et al., 2022). Regardless, all gymnosperms in our study were evergreen and generally leafed out later than deciduous species suggesting that accurately timing leaf out in early spring is less important since they can rely on previous year foliage initially (as long as they maintain the appropriate cold tolerance, discussed more below).

### Warming impacts on cold tolerance

Overall, cold tolerance was lost over the course of our study reflecting the natural dehardening process across the winter and spring seasons (Kalberer et al., 2006). Stems of deciduous species experienced relatively little change in cold tolerance when exposed to two weeks of warming. These results corroborate prior studies demonstrating that stems are less responsive than needles and leaf tissue (Li et al., 2002). Consistent with this finding, needles of most evergreen species have shown substantial reductions in cold tolerance in response to warming (Buchner & Neuner, 2011). For example, needles of *Juniperus virginiana, Picea rubens, Pinus strobus,* and *Pinus resinosa* had the largest reductions in cold tolerance during the first sampling event in March, bringing them closer to historical daily minimum temperatures and the potential for cold damage. During the second and third sampling event, both *pinus* species and *Abies balsamea* experienced major losses of cold tolerance in response to warming but maintained larger safety margins from historical cold temperatures. Although some research suggests that REL may underestimate pine cold tolerance in midwinter, our results still support the notion that many pines can tolerate colder temperatures than many other species in our study, but also that pines can de-acclimate rapidly in response to warming (Sutinen et al., 1992; Man et al., 2017). *Juniperus virginiana* and *Picea rubens* were generally the least cold tolerant species, aligning with *Juniperus virginiana*’s southern distribution and other work demonstrating that *Picea rubens* is only moderately cold tolerant and susceptible to winter injury (DeHayes et al., 2001). Combined, these results suggest that anomalously warm periods during winter, if followed by the return of extreme cold, may have significant impacts, particularly on needles of evergreen species in this region.

Timing leaf out correctly to avoid freezing damage is critical because damage can reduce carbon stores, cause biomass loss, limit growth, and drive mortality (Lenz et al., 2016; Reinmann et al., 2023; Muffler et al., 2024). While we observed little to no visible damage after seedlings experienced freezing temperatures following warming treatments, this could be due to our experiment taking place during a naturally relatively warm later-winter, where the damage risk from freezing temperatures was low. Across all species in our study, the majority of tissue damage from refreezing was below 10% per tree and the species that experience the most damage were *Betula papyrifera* (fast-growing, deciduous pioneer species) and *Juniperus virginiana* (beyond its northern range at our study site). So, while these results suggest that likelihood of refreezing damage is related to life history strategy and location relative to native range, damage never caused mortality or had an affect on height or diameter growth the following season. Additionally, our warming treatments did not affect the height and diameter of seedlings contrasting the widespread notion that longer growing seasons leads to more growth (Keenan et al., 2014; Grossiord et al., 2022). Growth responses may not have been detectable in our relatively short study and future work could examine the potentially different effects of extended growing seasons compared to warmer overall growing conditions (Reich et al., 2015).

### Conclusion

Our study highlights that understanding the links between environmental cues and phenological sensitivities is important in determining the future of spring phenology with climate change.

Although we generally found that longer warming events have a greater impact on leafing phenology, even short warming periods in mid-winter caused an average 4.3 day advancement of leaf out. Furthermore, the timing of leaf out and loss of cold tolerance depends on the time of year and length of warming experienced.

These results are particularly important for mixedwood forests of northeastern U.S. where diverse assemblages of leaf functional types (deciduous vs. evergreen) and life history strategies intersect at a regional ecotone where many species are at their southern or northern range margin. Our results suggest that, if they are able to avoid cold damage to needles, winter warming events may provide opportunities for evergreen species to extend their growing season. In contrast, except for fast growing pioneers, deciduous species may be less responsive to winter warming events helping them avoid potential damage while still timing spring leaf out to capture early springs. Combined, these results suggest that the likelihood of exposure to extreme cold following winter warming events will be critical to determining the future of these forests.

## Acknowledgements

This research was funded by the Northeastern States Research Cooperative through funding made available by the USDA Forest Service, Northern Research Station. The conclusions and opinions in this paper are those of the authors and not of the NSRC, the Forest Service, or the USDA. This project was also part of McIntire Stennis Project Number ME0-42121 and ME0-22502 administered through the Maine Agricultural and Forest Experiment Station. We would like to thank Alexandra Barry, Peter Breigenzer, Paige Cormier, Megan Grega, Cassie LaCasse, Emily MacDonald, and Brigid Mrenna for help with planting and data collection. We also want to acknowledge Brad Libby for facilitating this research at the Roger Clapp Greenhouse Complex.

## Author contributions

Laura Pinover, Nicholas Fisichelli, Nicole Rogers, Yong-Jiang Zhang, and Jay Wason designed the study and experiment. Laura Pinover, John Butnor, Paula Murakami, and Jay Wason collected and analyzed data. Laura Pinover and Jay Wason wrote the manuscript with contributions from all authors.

## Data availability

All data used in the production of this manuscript will be made publicly available via Dryad upon acceptance of the article for publication.

## Supporting Information

**Table S1** Accumulated growing degree days (GDD) for each cold tolerance sample.

**Fig. S1** Daily average temperatures throughout the duration of the study from inside the greenhouse and outside.

**Fig. S2** Examples of budburst and leaf out phenophases from study species.

**Fig. S3** Pair plot of fixed effects; Growing Degree Days, Chilling Days, and Photoperiod.

**Fig. S4** Relative height growth of study species from the 2024 growing season.

**Fig. S5** Relative height growth of study species from treatments.

**Fig. S6** Relative diameter growth of study species from the 2024 growing season.

**Fig. S7** Relative diameter growth of study species from treatments.

**Fig. S8** Day of year when budburst appeared for control study species.

**Fig. S9** Difference between treatment budburst date from control budburst date.

**Fig. S10** Difference in budburst day of year among each species and each warming treatment and control.

**Fig. S11** Effect sizes for budburst predictors (Growing Degree Days, photoperiod, and the interaction between GDD and photoperiod for all study species.

**Fig. S12** Effect sizes for budburst predictors (Growing Degree Days, Chilling Days, and the interaction between GDD and Chilling Days for all study species.

## References

Abramoff, R. Z., & Finzi, A. C. (2015). Are above-and below-ground phenology in sync? New Phytologist, 205(3), 1054–1061. 10.1111/nph.13111

Asse, D., Chuine, I., Vitasse, Y., Yoccoz, N. G., Delpierre, N., Badeau, V., Delestrade, A., & Randin, C. F. (2018). Warmer winters reduce the advance of tree spring phenology induced by warmer springs in the Alps. Agricultural and Forest Meteorology, 252, 220–230. 10.1016/j.agrformet.2018.01.030

Baumgarten, F., Gessler, A., & Vitasse, Y. (2023). No risk—no fun: Penalty and recovery from spring frost damage in deciduous temperate trees. Functional Ecology, 37(3), 648–663. 10.1111/1365-2435.14243

Bhattacharya, A. (2022). Effect of Low Temperature Stress on Photosynthesis and Allied Traits: A Review. In A. Bhattacharya (Ed.), Physiological Processes in Plants Under Low Temperature Stress (pp. 199–297). Springer. 10.1007/978-981-16-9037-2_3

Buchner, O., & Neuner, G. (2011). Winter frost resistance of Pinus cembra measured in situ at the alpine timberline as affected by temperature conditions. Tree Physiology, 31(11), 1217–1227. 10.1093/treephys/tpr103

Butnor, J. R., Wilson, C. P., Bakır, M., D’Amato, A. W., Flower, C. E., Hansen, C. F., Keller, S. R., Knight, K. S., & Murakami, P. F. (2024). Cold Tolerance Assay Reveals Evidence of Climate Adaptation Among American Elm (Ulmus americana L.) Genotypes. Forests, 15(11), Article 11. 10.3390/f15111843

Cannell, M. G. R., & Smith, R. I. (1983). Thermal Time, Chill Days and Prediction of Budburst in Picea sitchensis. Journal of Applied Ecology, 20(3), 951–963. 10.2307/2403139

Chang, C. Y.-Y., Bräutigam, K., Hüner, N. P. A., & Ensminger, I. (2021). Champions of winter survival: Cold acclimation and molecular regulation of cold hardiness in evergreen conifers. New Phytologist, 229(2), 675–691. 10.1111/nph.16904

DeHayes, D. H., Schaberg, P. G., & Strimbeck, G. R. (2001). Red Spruce (Picea rubens Sarg.) Cold Hardiness and Freezing Injury Susceptibility. In F. J. Bigras & S. J. Colombo (Eds.), Conifer Cold Hardiness (pp. 495–529). Springer Netherlands. 10.1007/978-94-015-9650-3_18

Ettinger, A. K., Chamberlain, C. J., Morales-Castilla, I., Buonaiuto, D. M., Flynn, D. F. B., Savas, T., Samaha, J. A., & Wolkovich, E. M. (2020). Winter temperatures predominate in spring phenological responses to warming. Nature Climate Change, 10(12), 1137– 1142. 10.1038/s41558-020-00917-3

Fahey, R. T. (2016). Variation in responsiveness of woody plant leaf out phenology to anomalous spring onset. Ecosphere, 7(2), e01209. 10.1002/ecs2.1209

Fenner, M. (1987). Seedlings. New Phytologist, 106(s1), 35–47. 10.1111/j.1469-8137.1987.tb04681.x

Flynn, D. F. B., & Wolkovich, E. M. (2018). Temperature and photoperiod drive spring phenology across all species in a temperate forest community. New Phytologist, 219(4), 1353–1362. 10.1111/nph.15232

Frelich, L. E., Montgomery, R. A., & Reich, P. B. (2021). Seven ways a warming climate can kill the southern boreal forest. Forests, 12(5). 10.3390/f12050560

Grossiord, C., Bachofen, C., Gisler, J., Mas, E., Vitasse, Y., & Didion-Gency, M. (2022). Warming may extend tree growing seasons and compensate for reduced carbon uptake during dry periods. Journal of Ecology, 110(7), 1575–1589. 10.1111/1365-2745.13892

Kalberer, S. R., Wisniewski, M., & Arora, R. (2006). Deacclimation and reacclimation of cold-hardy plants: Current understanding and emerging concepts. Plant Science, 171(1), 3–16. 10.1016/j.plantsci.2006.02.013

Keenan, T. F., Gray, J., Friedl, M. A., Toomey, M., Bohrer, G., Hollinger, D. Y., Munger, J. W., O’Keefe, J., Schmid, H. P., Wing, I. S., Yang, B., & Richardson, A. D. (2014). Net carbon uptake has increased through warming-induced changes in temperate forest phenology. Nature Climate Change, 4(7), 598–604. 10.1038/nclimate2253

Körner, C., & Basler, D. (2010). Phenology Under Global Warming. Science, 327(5972), 1461–1462. 10.1126/science.1186473

Körner, C., Basler, D., Hoch, G., Kollas, C., Lenz, A., Randin, C. F., Vitasse, Y., & Zimmermann, N. E. (2016). Where, why and how? Explaining the low-temperature range limits of temperate tree species. Journal of Ecology, 104(4), 1076–1088. 10.1111/1365-2745.12574

Kreyling, J., Schmid, S., & Aas, G. (2015). Cold tolerance of tree species is related to the climate of their native ranges. Journal of Biogeography, 42(1), 156–166. 10.1111/jbi.12411

Ladwig, L. M., Chandler, J. L., Guiden, P. W., & Henn, J. J. (2019). Extreme winter warm event causes exceptionally early bud break for many woody species. Ecosphere, 10(1), e02542. 10.1002/ecs2.2542

Lenz, A., Hoch, G., Körner, C., & Vitasse, Y. (2016). Convergence of leaf-out towards minimum risk of freezing damage in temperate trees. Functional Ecology, 30(9), 1480–1490. 10.1111/1365-2435.12623

Li, C., Puhakainen, T., Welling, A., Viherä-Aarnio, A., Ernstsen, A., Junttila, O., Heino, P., & Palva, E. T. (2002). Cold acclimation in silver birch (Betula pendula). Development of freezing tolerance in different tissues and climatic ecotypes. Physiologia Plantarum, 116(4), 478–488. 10.1034/j.1399-3054.2002.1160406.x

Liu, H., Wang, H., Li, N., Shao, J., Zhou, X., van Groenigen, K. J., & Thakur, M. P. (2022). Phenological mismatches between above-and belowground plant responses to climate warming. Nature Climate Change, 12(1), 97–102. 10.1038/s41558-021-01244-x

Lu, C., van Groenigen, K. J., Gillespie, M. A. K., Hollister, R. D., Post, E., Cooper, E. J., Welker, J. M., Huang, Y., Min, X., Chen, J., Jónsdóttir, I. S., Mauritz, M., Cannone, N., Natali, S. M., Schuur, E., Molau, U., Yan, T., Wang, H., He, J.-S., & Liu, H. (2025). Diminishing warming effects on plant phenology over time. New Phytologist, 245(2), 523–533. 10.1111/nph.20019

Malyshev, A. V., Blume-Werry, G., Spiller, O., Smiljanić, M., Weigel, R., Kolb, A., Nze, B. Y., Märker, F., Sommer, F. C.-F. J., Kinley, K., Ziegler, J., Pasang, P., Mahara, R., Joshi, S., Heinsohn, V., & Kreyling, J. (2023). Warming nondormant tree roots advances aboveground spring phenology in temperate trees. New Phytologist, 240(6), 2276–2287. 10.1111/nph.19304

Man, R., Colombo, S., Lu, P., & Dang, Q.-L. (2016). Effects of winter warming on cold hardiness and spring budbreak of four boreal conifers. Botany, 94(2), 117–126. 10.1139/cjb-2015-0181

Man, R., Colombo, S., Lu, P., Li, J., & Dang, Q.-L. (2014). Trembling aspen, balsam poplar, and white birch respond differently to experimental warming in winter months. Canadian Journal of Forest Research, 44(12), 1469–1476. 10.1139/cjfr-2014-0302

Man, R., Lu, P., & Dang, Q.-L. (2017). Cold hardiness of white spruce, black spruce, jack pine, and lodgepole pine needles during dehardening. Canadian Journal of Forest Research, 47(8), 1116–1122. 10.1139/cjfr-2017-0119

Marquis, B., & Lajoie, G. (2023). Experimental exposure to winter thaws reveals tipping point in yellow birch bud mortality and phenology in the northern temperate forest of Québec, Canada (p. 2023.10.20.563331). bioRxiv. 10.1101/2023.10.20.563331

Marquis, B., & Lajoie, G. (2024). Experimental exposure to winter thaws reveals tipping point in yellow birch bud mortality and phenology in the northern temperate forest of Québec, Canada. Climate Change Ecology, 7, 100087. 10.1016/j.ecochg.2024.100087

Meng, L., Zhou, Y., Gu, L., Richardson, A. D., Peñuelas, J., Fu, Y., Wang, Y., Asrar, G. R., De Boeck, H. J., Mao, J., Zhang, Y., & Wang, Z. (2021). Photoperiod decelerates the advance of spring phenology of six deciduous tree species under climate warming. Global Change Biology, 27(12), 2914–2927. 10.1111/gcb.15575

Michelot, A., Simard, S., Rathgeber, C., Dufrêne, E., & Damesin, C. (2012). Comparing the intra-annual wood formation of three European species (Fagus sylvatica, Quercus petraea and Pinus sylvestris) as related to leaf phenology and non-structural carbohydrate dynamics. Tree Physiology, 32(8), 1033–1045. 10.1093/treephys/tps052

Miller-Rushing, A. J., Inouye, D. W., & Primack, R. B. (2008). How well do first flowering dates measure plant responses to climate change? The effects of population size and sampling frequency. Journal of Ecology, 96(6), 1289–1296. 10.1111/j.1365-2745.2008.01436.x

Montgomery, R. A., Rice, K. E., Stefanski, A., Rich, R. L., & Reich, P. B. (2020). Phenological responses of temperate and boreal trees to warming depend on ambient spring temperatures, leaf habit, and geographic range. Proceedings of the National Academy of Sciences of the United States of America, 117(19), 10397–10405. 10.1073/pnas.1917508117

Morin, X., Roy, J., Sonié, L., & Chuine, I. (2010). Changes in leaf phenology of three European oak species in response to experimental climate change. New Phytologist, 186(4), 900– 910. 10.1111/j.1469-8137.2010.03252.x

Muffler, L., Weigel, R., Beil, I., Leuschner, C., Schmeddes, J., & Kreyling, J. (2024). Winter and spring frost events delay leaf-out, hamper growth and increase mortality in European beech seedlings, with weaker effects of subsequent frosts. Ecology and Evolution, 14(7), e70028. 10.1002/ece3.70028

Nanninga, C., Buyarski, C. R., Pretorius, A. M., & Montgomery, R. A. (2017). Increased exposure to chilling advances the time to budburst in North American tree species. Tree Physiology, 37(12), 1727–1738. 10.1093/treephys/tpx136

Pan, Y.-Q., Zeng, X., Chen, W.-D., Tang, X.-R., Dai, K., Du, Y.-J., & Song, X.-Q. (2022). Chilling rather than photoperiod controls budburst for gymnosperm species in subtropical China. Journal of Plant Ecology, 15(1), 100–110. 10.1093/jpe/rtab076

Panchen, Z. A., Primack, R. B., Nordt, B., Ellwood, E. R., Stevens, A.-D., Renner, S. S., Willis, C. G., Fahey, R., Whittemore, A., Du, Y., & Davis, C. C. (2014). Leaf out times of temperate woody plants are related to phylogeny, deciduousness, growth habit and wood anatomy. New Phytologist, 203(4), 1208–1219. 10.1111/nph.12892

Parmesan, C. (2007). Influences of species, latitudes and methodologies on estimates of phenological response to global warming. Global Change Biology, 13(9), 1860–1872. 10.1111/j.1365-2486.2007.01404.x

Posit team. (2024). *RStudio: Integrated Development Environment for R* [Computer software]. Posit Software, PBC. http://www.posit.co/

R Core Team. (2024). *R: A Language and Environment for Statistical Computing* [Computer software]. R Foundation for Statistical Computing. https://www.R-project.org/

Reich, P. B., Sendall, K. M., Rice, K., Rich, R. L., Stefanski, A., Hobbie, S. E., & Montgomery, R. A. (2015). Geographic range predicts photosynthetic and growth response to warming in co-occurring tree species. Nature Climate Change, 5(2), 148–152. 10.1038/nclimate2497

Reinmann, A. B., Bowers, J. T., Kaur, P., & Kohler, C. (2023). Compensatory responses of leaf physiology reduce effects of spring frost defoliation on temperate forest tree carbon uptake. Frontiers in Forests and Global Change, 6. 10.3389/ffgc.2023.988233

Richardson, A. D., Andy Black, T., Ciais, P., Delbart, N., Friedl, M. A., Gobron, N., Hollinger, D. Y., Kutsch, W. L., Longdoz, B., Luyssaert, S., Migliavacca, M., Montagnani, L., William Munger, J., Moors, E., Piao, S., Rebmann, C., Reichstein, M., Saigusa, N., Tomelleri, E.,…Varlagin, A. (2010). Influence of spring and autumn phenological transitions on forest ecosystem productivity. Philosophical Transactions of the Royal Society B: Biological Sciences, 365(1555), 3227–3246. 10.1098/rstb.2010.0102

Richardson, A. D., Keenan, T. F., Migliavacca, M., Ryu, Y., Sonnentag, O., & Toomey, M. (2013). Climate change, phenology, and phenological control of vegetation feedbacks to the climate system. Agricultural and Forest Meteorology, 169, 156–173. 10.1016/j.agrformet.2012.09.012

Ritz, C., Baty, F., Streibig, J. C., & Gerhard, D. (2015). Dose-Response Analysis Using R. PLOS ONE, 10(12), e0146021. 10.1371/journal.pone.0146021

Rosemartin, A. H., Denny, E. G., Gerst, K. L., Marsh, R. L., Posthumus, E. E., Crimmins, T. M., & Weltzin, J. (2018). USA National Phenology Network observational data documentation. In Open-File Report (2018–1060). U.S. Geological Survey. 10.3133/ofr20181060

Satake, A., Nagahama, A., & Sasaki, E. (2022). A cross-scale approach to unravel the molecular basis of plant phenology in temperate and tropical climates. New Phytologist, 233(6), 2340–2353. 10.1111/nph.17897

Savage, J. A., Kiecker, T., McMann, N., Park, D., Rothendler, M., & Mosher, K. (2022). Leaf out time correlates with wood anatomy across large geographic scales and within local communities. New Phytologist, 235(3), 953–964. 10.1111/nph.18041

Schaberg, P. G., Lazarus, B. E., Hawley, G. J., Halman, J. M., Borer, C. H., & Hansen, C. F. (2011). Assessment of weather-associated causes of red spruce winter injury and consequences to aboveground carbon sequestration. Canadian Journal of Forest Research, 41(2), 359–369. 10.1139/X10-202

Skinner, M., & Parker, B. (1994). Field Guide for Monitoring Sugar Maple Bud Development – Maple Research. https://mapleresearch.org/pub/maplebudfieldguide-2/

Strimbeck, G. R., Kjellsen, T. D., Schaberg, P. G., & Murakami, P. F. (2007). Cold in the common garden: Comparative low-temperature tolerance of boreal and temperate conifer foliage. Trees, 21(5), 557–567. 10.1007/s00468-007-0151-1

Stuble, K. L., Bennion, L. D., & Kuebbing, S. E. (2021). Plant phenological responses to experimental warming—A synthesis. Global Change Biology, 27(17), 4110–4124. 10.1111/gcb.15685

Sutinen, M.-L., Palta, J. P., & Reich, P. B. (1992). Seasonal differences in freezing stress resistance of needles of Pinus nigra and Pinus resinosa: Evaluation of the electrolyte leakage method. Tree Physiology, 11(3), 241–254. 10.1093/treephys/11.3.241

Thackeray, S. J., Henrys, P. A., Hemming, D., Bell, J. R., Botham, M. S., Burthe, S., Helaouet, P., Johns, D. G., Jones, I. D., Leech, D. I., Mackay, E. B., Massimino, D., Atkinson, S., Bacon, P. J., Brereton, T. M., Carvalho, L., Clutton-Brock, T. H., Duck, C., Edwards, M.,…Wanless, S. (2016). Phenological sensitivity to climate across taxa and trophic levels. Nature, 535(7611), 241–245. 10.1038/nature18608

Thieurmel, B., & Elmarhraoui, A. (2022). *suncalc: Compute Sun Position, Sunlight Phases, Moon Position and Lunar Phase* (Version 0.5.1) [Computer software]. https://CRAN.R-project.org/package=suncalc

Tukey, J. W. (1953). Section of Mathematics and Engineering: Some Selected Quick and Easy Methods of Statistical Analysis. Transactions of the New York Academy of Sciences, 16(2 Series II), 88–97. 10.1111/j.2164-0947.1953.tb01326.x

Vitasse, Y., Lenz, A., Hoch, G., & Körner, C. (2014). Earlier leaf-out rather than difference in freezing resistance puts juvenile trees at greater risk of damage than adult trees. Journal of Ecology, 102(4), 981–988. 10.1111/1365-2745.12251

Winkler, D. E., Garbowski, M., Kožić, K., Ladouceur, E., Larson, J., Martin, S., Rosche, C., Roscher, C., Slate, M. L., & Korell, L. (2024). Facilitating comparable research in seedling functional ecology. Methods in Ecology and Evolution, 15(3), 464–476. 10.1111/2041-210X.14288

Zettlemoyer, M. A., Schultheis, E. H., & Lau, J. A. (2019). Phenology in a warming world: Differences between native and non-native plant species. Ecology Letters, 22(8), 1253– 1263. 10.1111/ele.13290

Zhang, H., Chuine, I., Regnier, P., Ciais, P., & Yuan, W. (2022). Deciphering the multiple effects of climate warming on the temporal shift of leaf unfolding. Nature Climate Change, 12(2), 193–199. 10.1038/s41558-021-01261-w

Zohner, C. M., Benito, B. M., Svenning, J.-C., & Renner, S. S. (2016). Day length unlikely to constrain climate-driven shifts in leaf-out times of northern woody plants. Nature Climate Change, 6(12), 1120–1123. 10.1038/nclimate3138

Zohner, C. M., & Renner, S. S. (2015). Perception of photoperiod in individual buds of mature trees regulates leaf-out. New Phytologist, 208(4), 1023–1030. 10.1111/nph.13510

