## Supplemental Information for "Earlier leaf out and loss of cold tolerance for northeastern U.S. trees in response to winter warming events and early springs"

### *New Phytologist* Supporting Information

Article acceptance date: Click here to enter a date.

The following Supporting Information is available for this article:

**Table S1** Accumulated Growing Degree Days (GDD) represented for each cold tolerance sampling date for both control and warmed treatments.

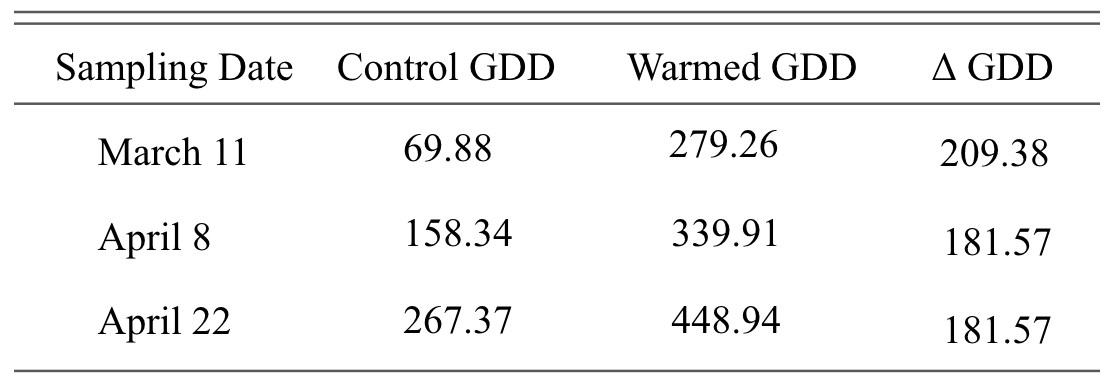

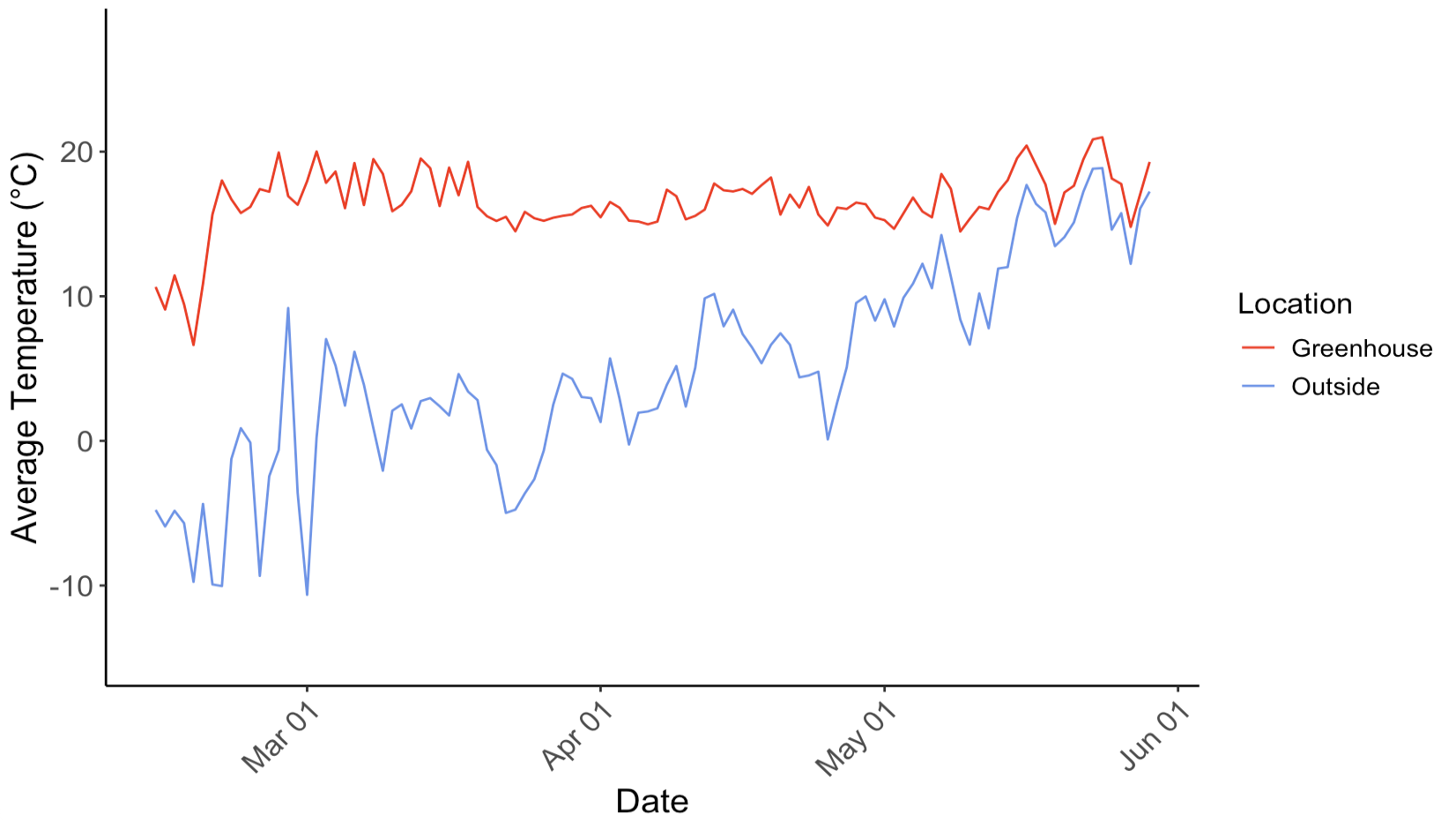

**Fig. S1** Daily average temperatures from the greenhouse and outside Onset HOBO External Temperature/RH Sensor Data Loggers. Two HOBO Data Loggers were used in the outside and greenhouse conditions. The average from the two corresponding to the environment was taken to get one daily average value. Dates included cover the duration of all warming treatments.

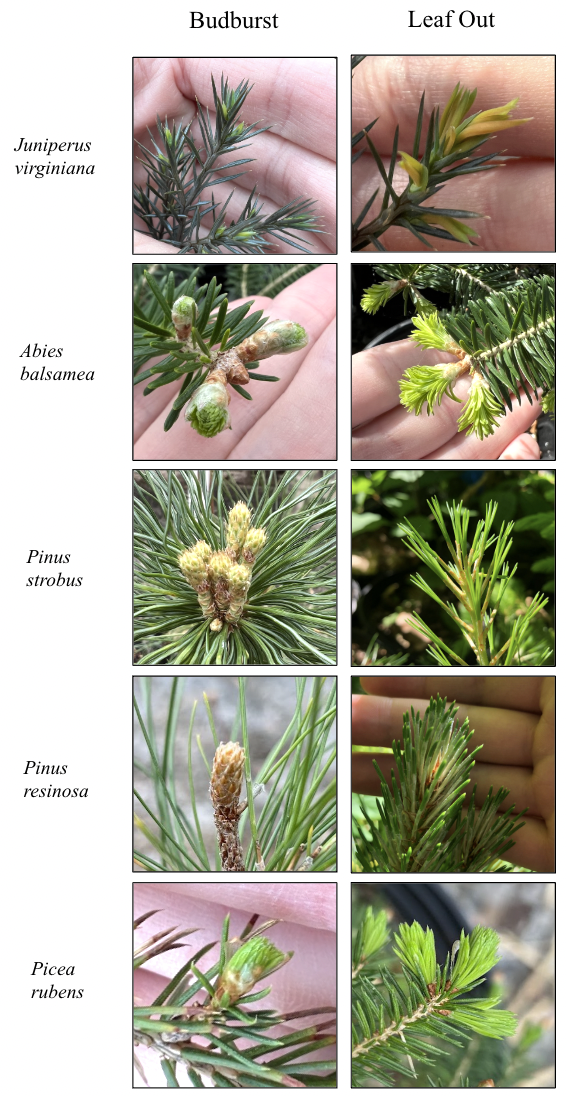

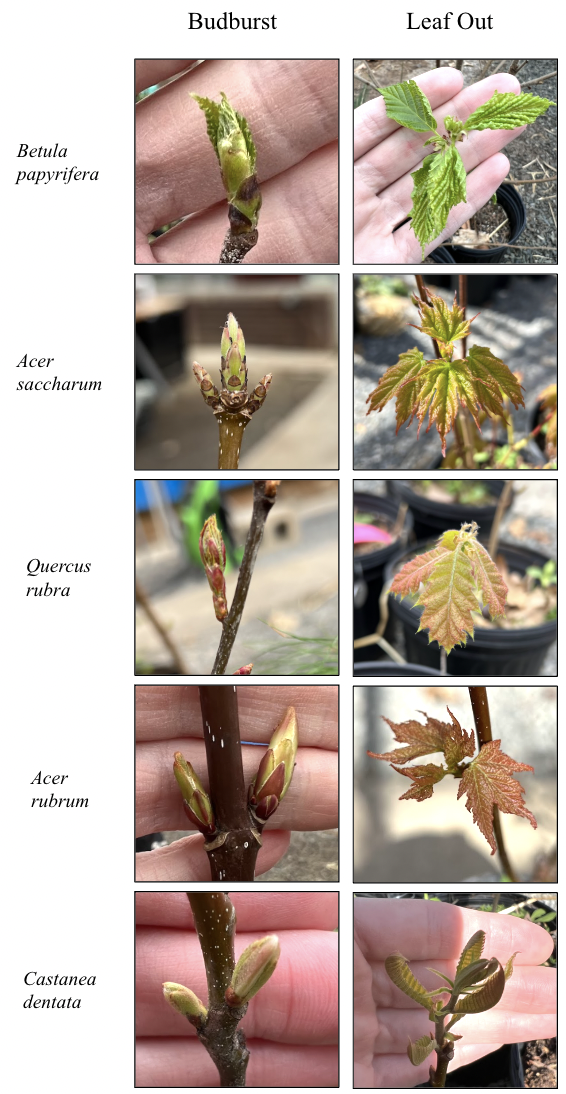

**Fig. S2** Examples of the budburst and leaf out phenophases from the phenology guides that we created. Every study species is represented in these examples.

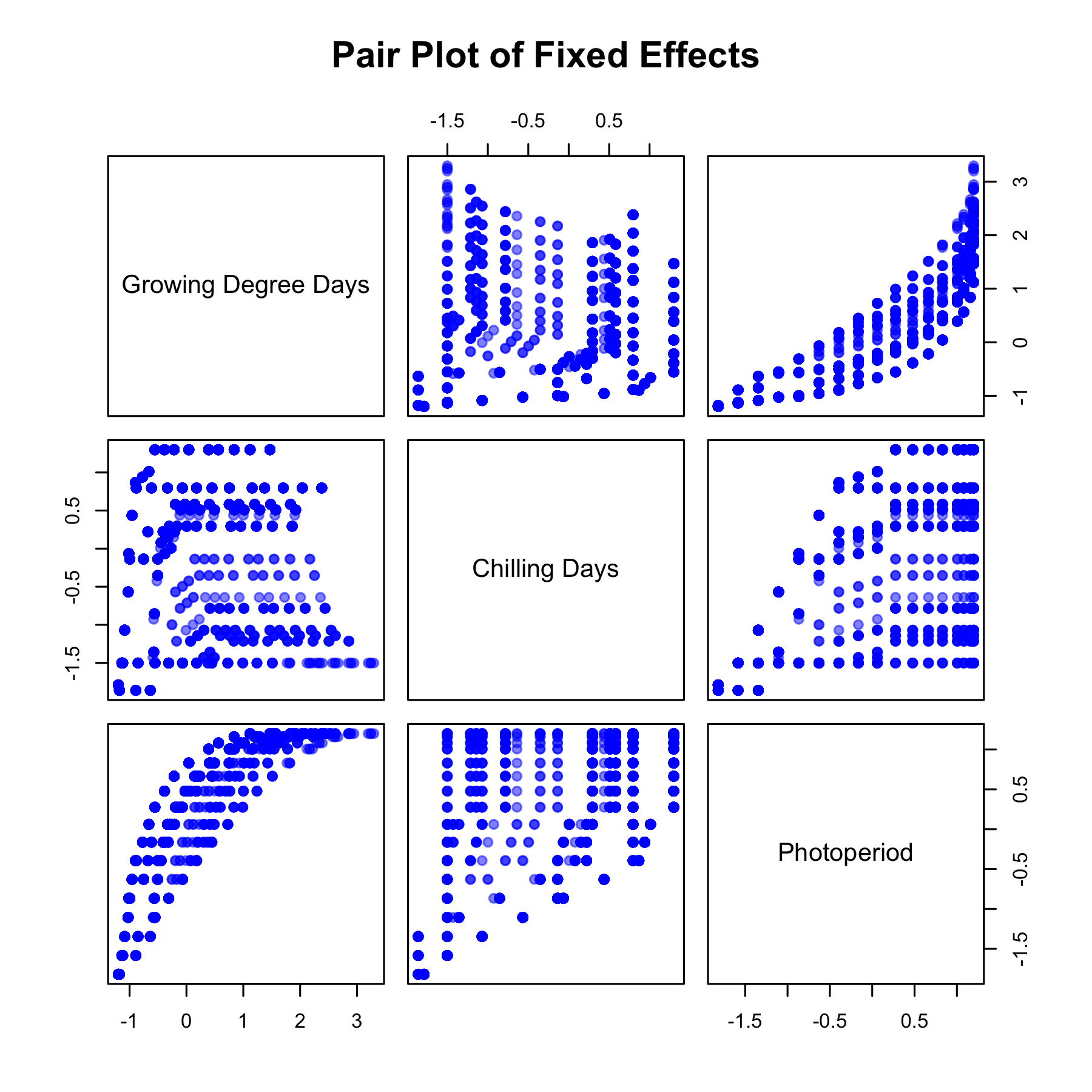

**Fig. S3** Pair plot of the fixed effects; Growing Degree Days, Chilling Days, and Photoperiod. All fixed effects are scaled.

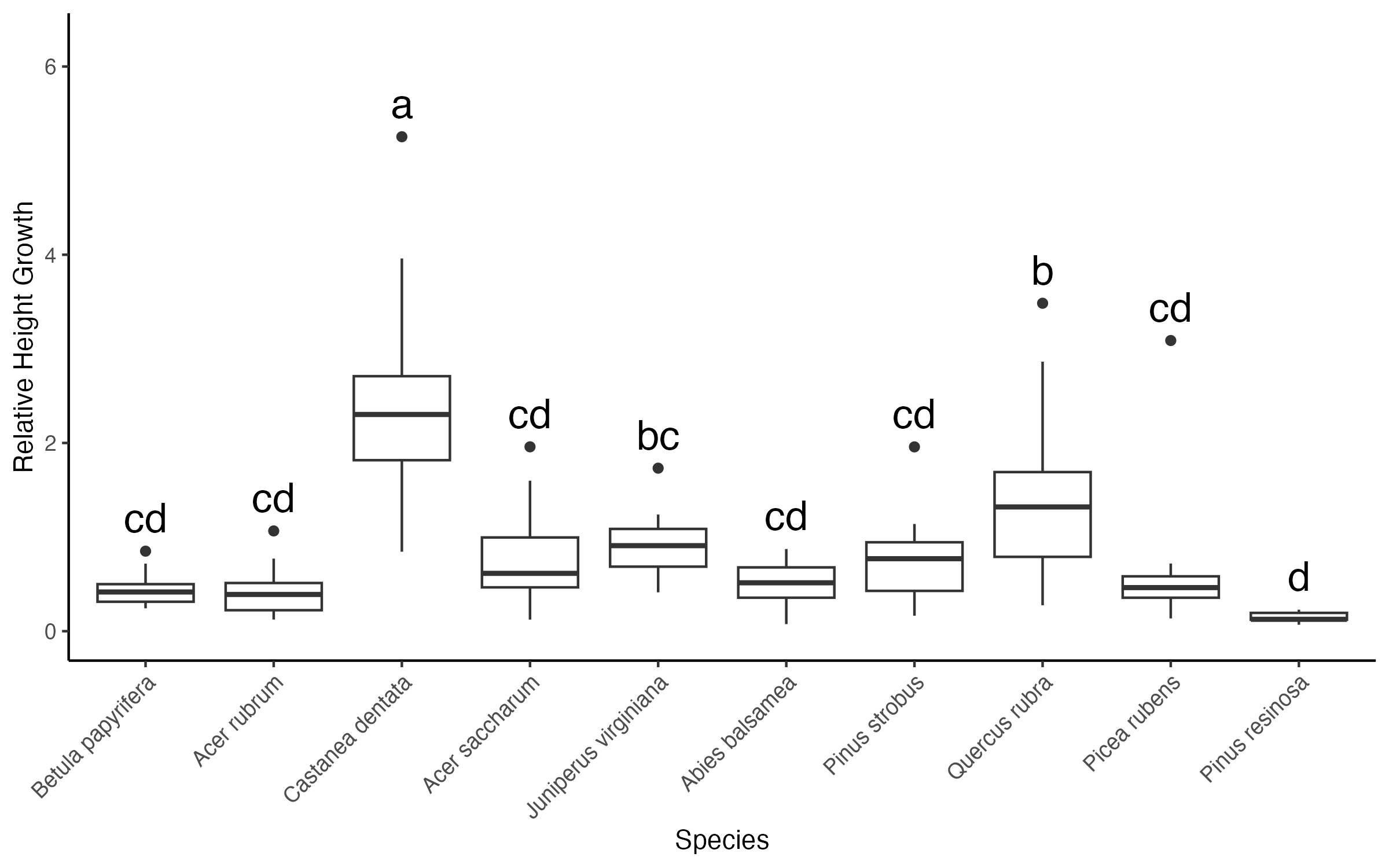

**Fig. S4** Relative height growth of study species from the 2024 growing season. Significance in height growth among species is indicated by different letters. Species are ordered by leaf out date, starting with the earliest to the latest.

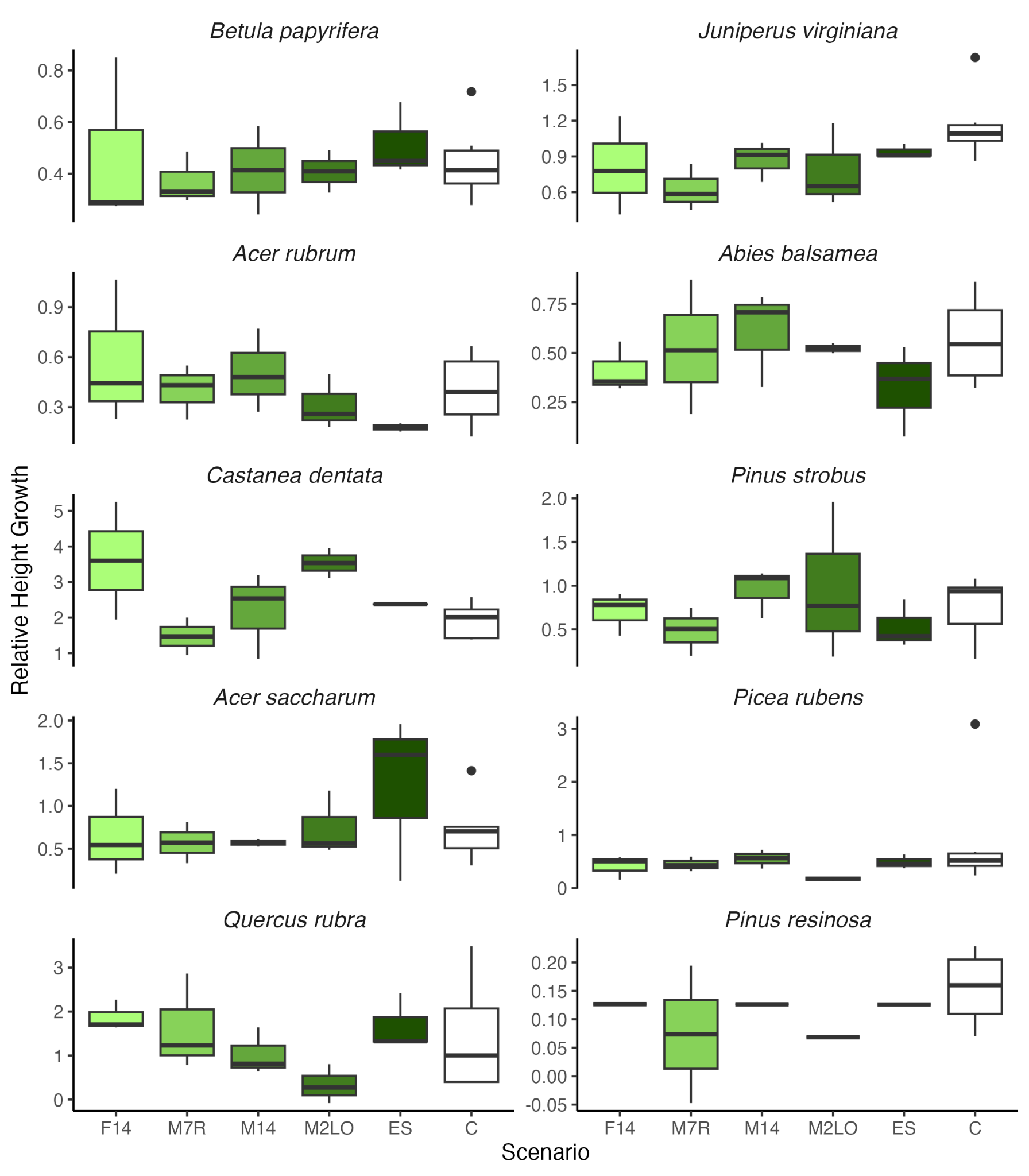

**Fig. S5** Relative height growth of study species from each treatment from the 2024 growing season. Scenarios did not have a significant effect on height growth (p>0.05).

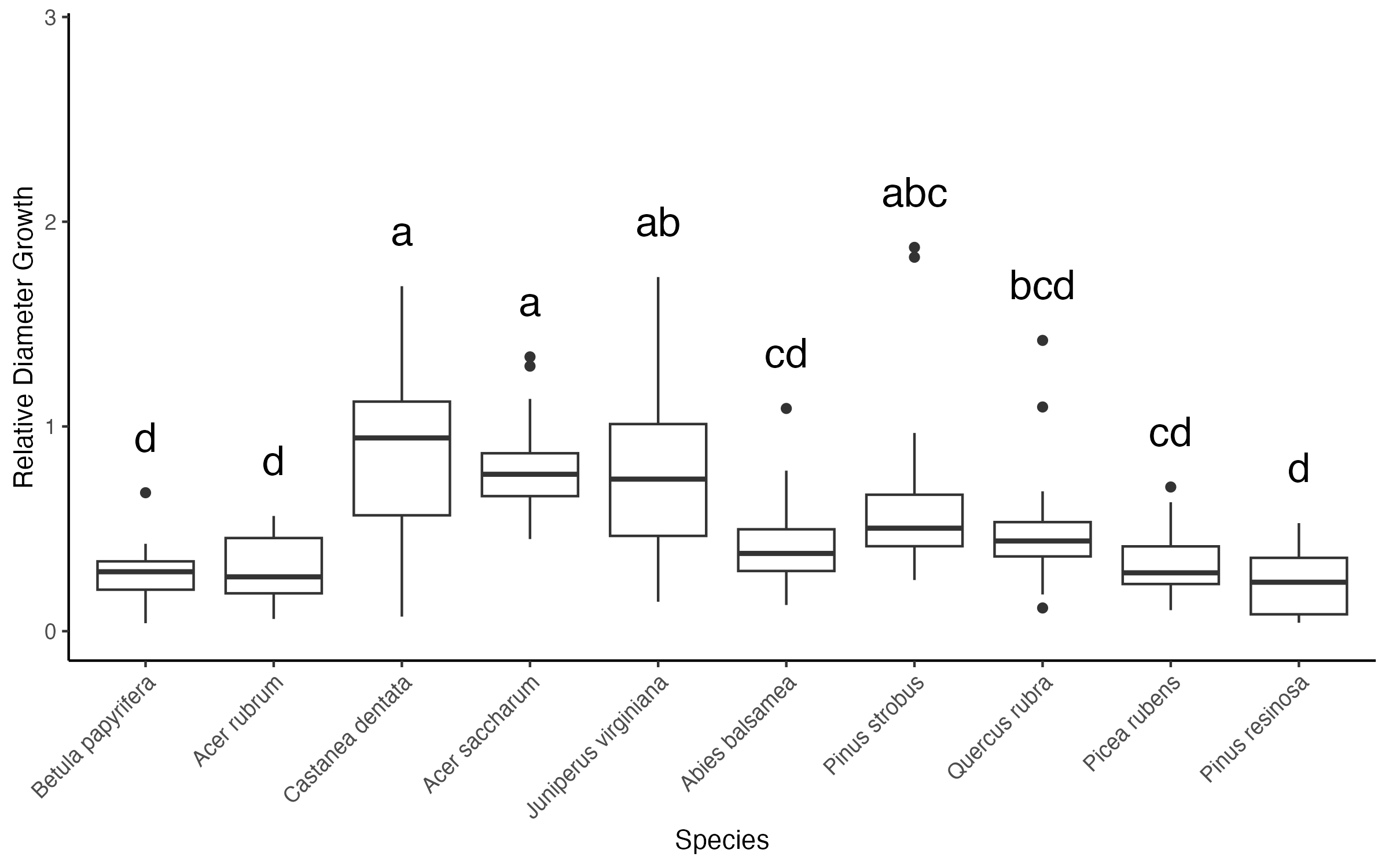

**Fig. S6** Relative diameter growth of study species from the 2024 growing season. Significance in height growth across species is indicated by different letters. Species are ordered by leaf out date, starting with the earliest to the latest.

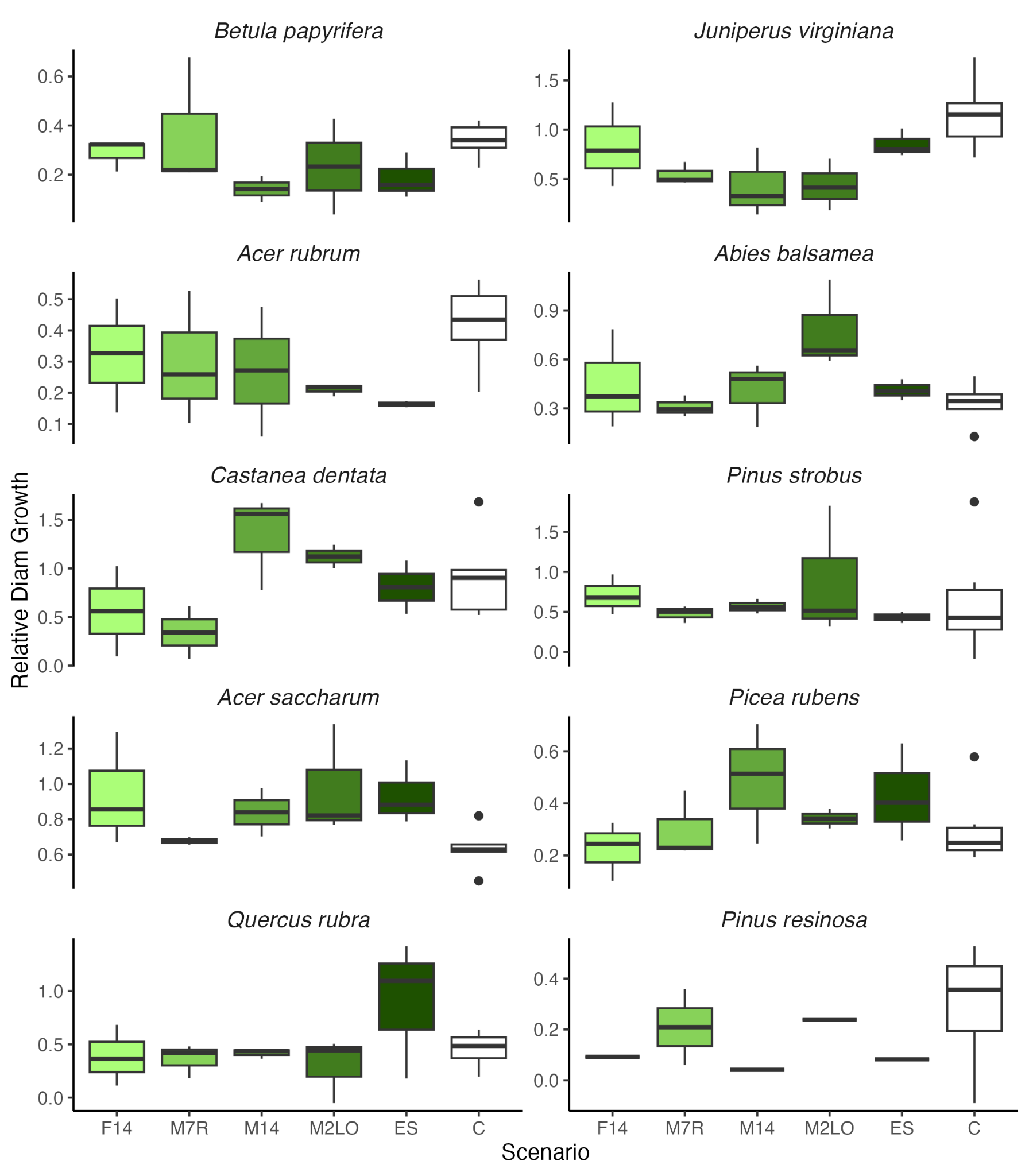

**Fig. S7** Relative diameter growth of study species from each scenario from the 2024 growing season. Scenarios did not have a significant effect on diameter growth (p>0.05).

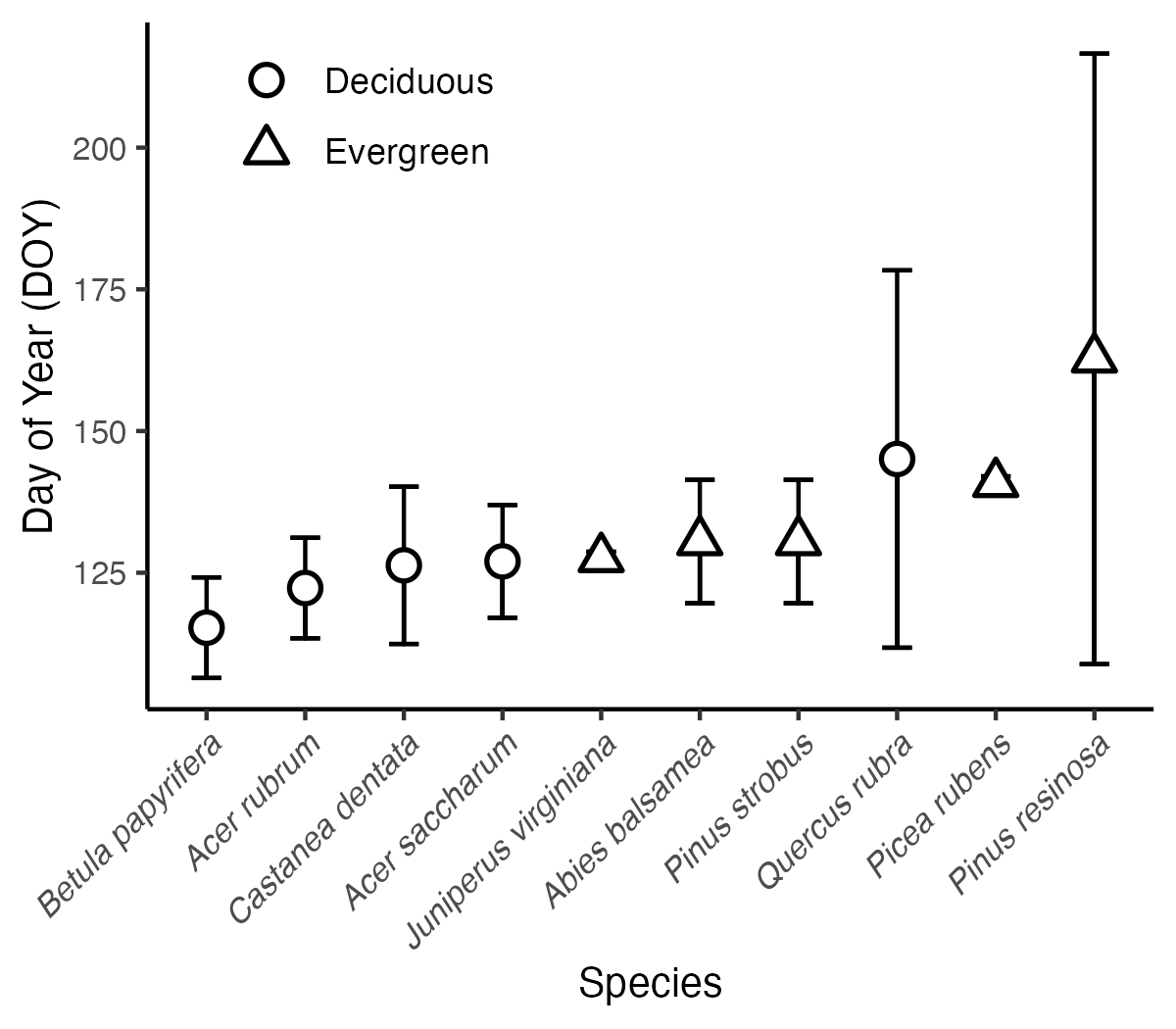

**Fig. S8** Day of year when majority budburst appeared for our control study species represented by 95% confidence intervals. Species with no visible error bars had little or no variation in budburst date that was detectable by our weekly resolution.

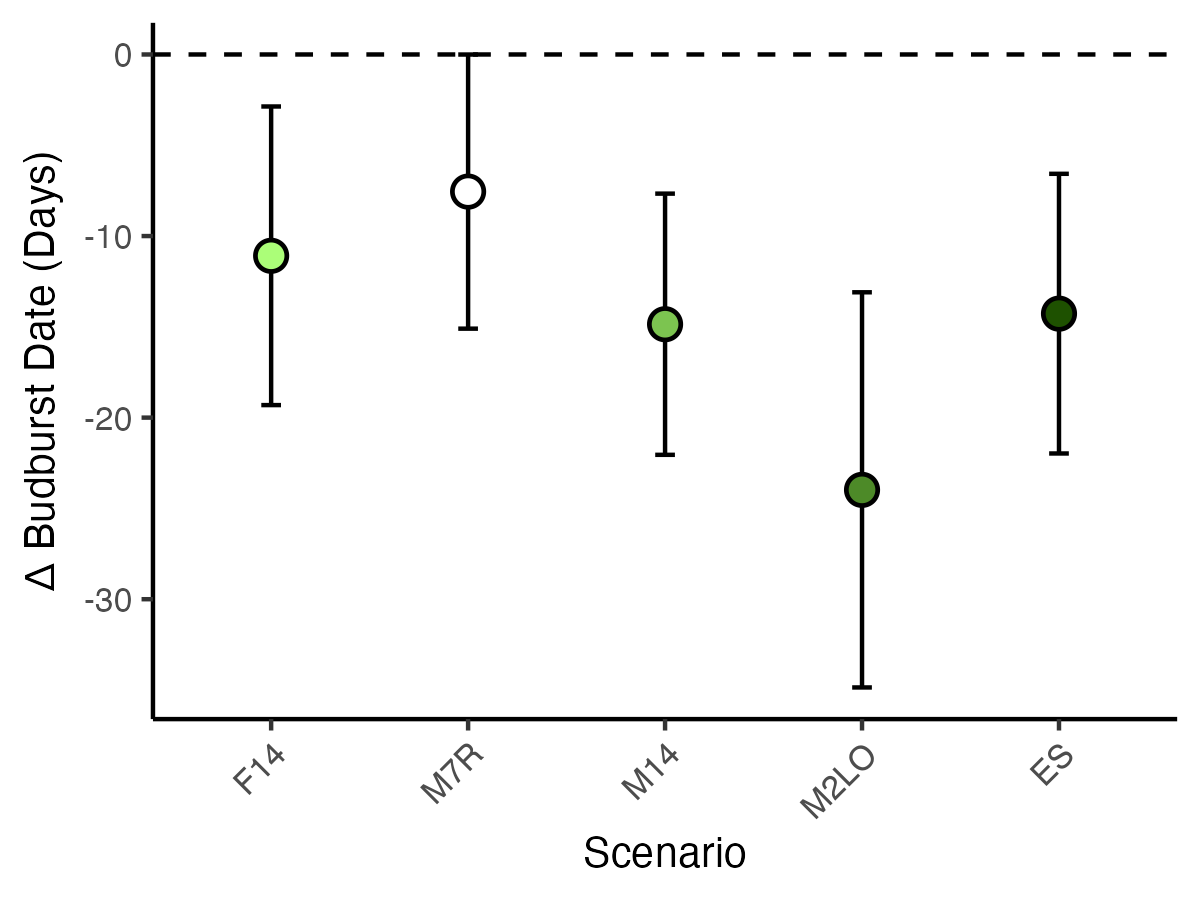

**Fig. S9** The mean difference between treatment budburst date from control budburst date represented by 95% confidence intervals. Significance among scenarios is accounted for when confidence intervals do not overlap.

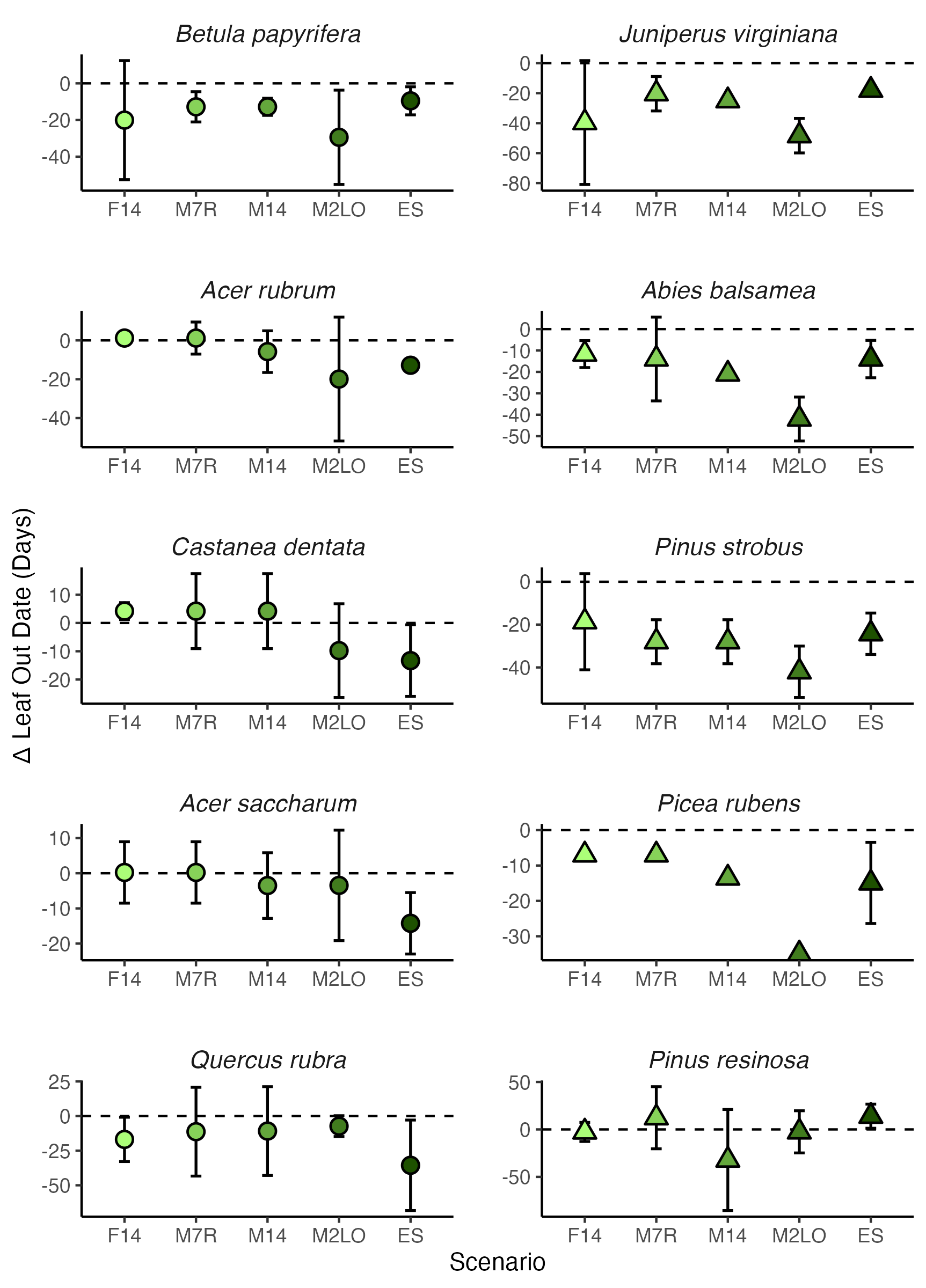

**Fig. S10** 95% confidence intervals for the difference in budburst day of year (DOY) among each species and each warming treatment and control. Each panel represents a species. Circles indicate deciduous species and triangles indicate evergreen species. Warming treatments are deemed significant if the 95% confidence intervals fall below 0. Species with each column are ordered by the budburst date for our controls, starting with the species to reach budburst earliest to the species to reach budburst the latest. Shape means with no visible error bars, have either error bars behind the shape or no variation in the mean

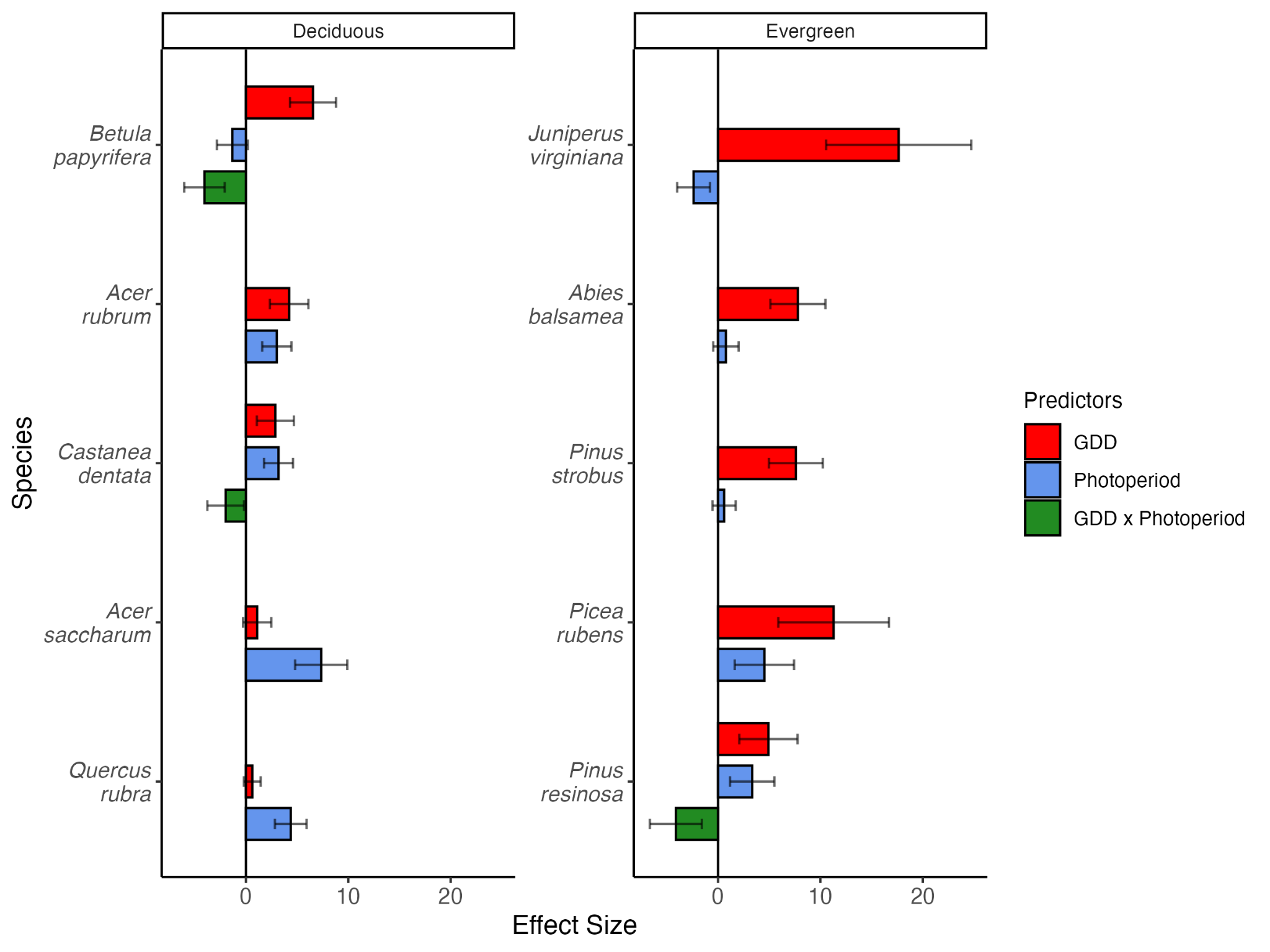

**Fig. S11** Effect sizes for budburst predictors (Growing Degree Days (GDD), photoperiod, and the interaction between GDD and photoperiod for all study species from generalized linear mixed effects models. Error bars are 95% confidence intervals. Individual models were created for each species. Columns are split by deciduous and evergreen species. Species with all three predictors present included the interaction between GDD and photoperiod in the model, whereas species with two predictors had the interaction removed from their models.

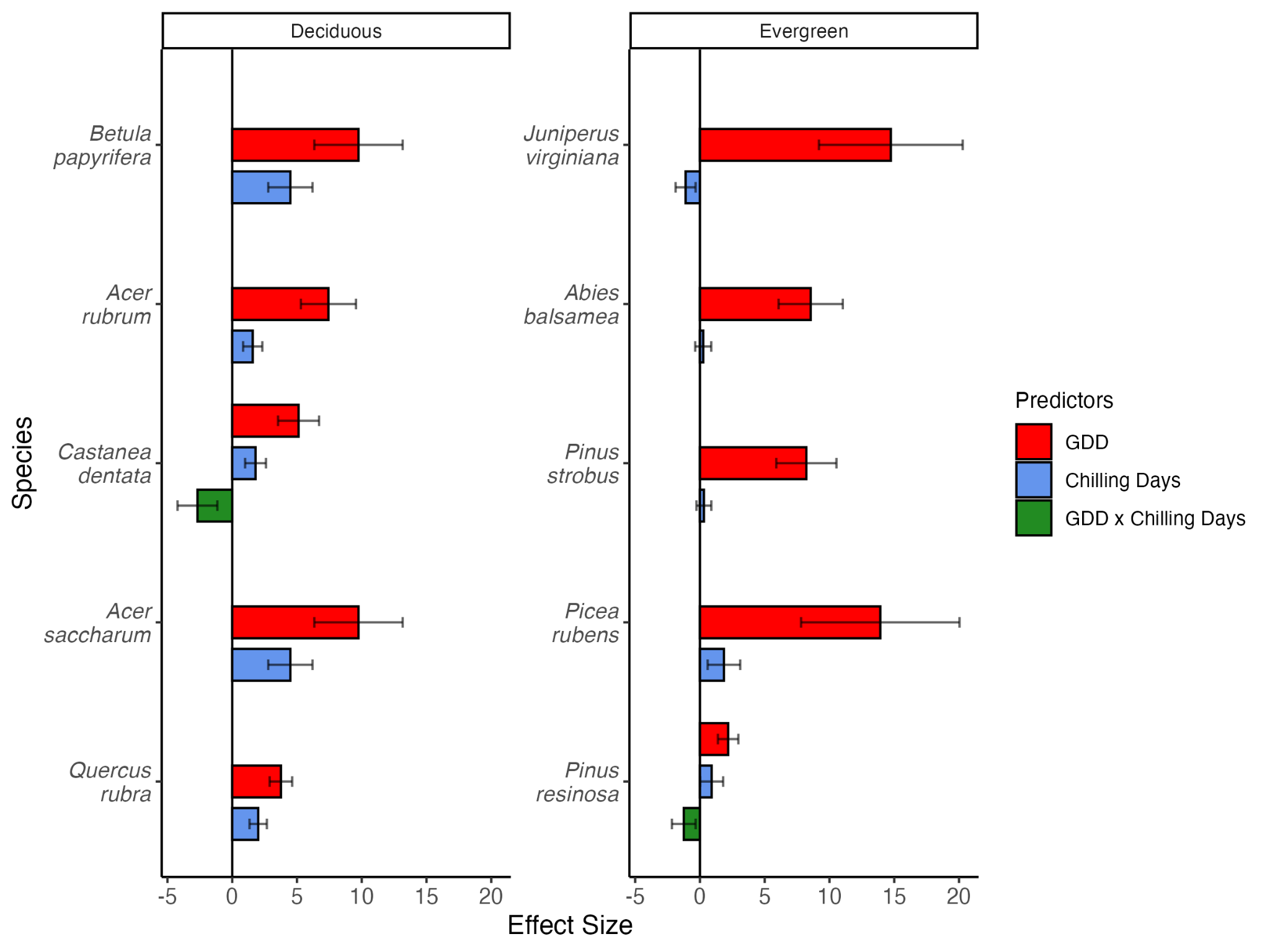

**Fig. S12** Effect sizes for budburst predictors (Growing Degree Days (GDD), Chilling Days, and the interaction between GDD and Chilling Days for all study species from generalized linear mixed effects models. Error bars are 95% confidence intervals. Individual models were created for each species. Columns are split by deciduous and evergreen species. Species with all three predictors present included the interaction between GDD and Chilling Days in the model, whereas species with two predictors had the interaction removed from their models.
